# Murine osteosarcoma recapitulates the driver landscape and genomic complexity of osteosarcoma evolution in humans

**DOI:** 10.64898/2026.04.27.721100

**Authors:** Geoffrey A. Smith, Ianthe A.E.M van Belzen, Mathieu Epinette, Emilia Herdes, Kim L. Mercer, Charlotte G. Butterworth, Alistair G. Rust, Adrienne M. Flanagan, Matthew G. Jones, Isidro Cortes-Ciriano, Tyler Jacks

## Abstract

Osteosarcoma (OS) genomes are characterized by complex genomic rearrangements (CGRs) that drive genomic instability and clonal diversification early in tumor evolution. As a result, OS tumors display high inter-patient variability, which has hindered molecular stratification and targeted therapeutic development. To study genomic complexity in OS and credential a genetically engineered mouse model of the disease (*Sp7-Cre Trp53*^*fl*^ Rb1^fl^), we performed high-depth and multi-region whole genome sequencing (WGS) of 35 tumor samples from 24 mice. Similar to human OS, the murine OS tumors (mOS) had a high number of somatic structural variants (158 per tumor) with low tumor mutational burden of single nucleotide variants (0.87 mutations/MB). CGRs were identified in 63% (15/24) of mOS cases, most frequently affecting chromosome 15 (33%, 8/24 mice) and resulting in *Myc* amplification in 6 mice, ranging from 5 to 104 copies. *Myc* amplification was verified with DNA FISH, long-read sequencing and gene expression data, which revealed examples of *Myc* amplification in both extrachromosomal circular DNA (ecDNA) and in derivative chromosomes generated by CGRs. *PTEN* loss occurred frequently (59% 12/22 mice), and contributed to osteosarcomagenesis, as demonstrated by tumor initiation with *in vivo* CRISPR/Cas9-mediated deletion experiments (2 mice). Together, these results demonstrate that a preclinical model of osteosarcoma can generate the genomic heterogeneity and complexity of the human disease, thereby facilitating research into mechanisms of tumor initiation and drivers of progression and relapse.

## Introduction

Osteosarcoma (OS) is the most common primary bone cancer and primarily affects adolescents and young adults.^1^ Treatment has not changed significantly in over 40 years. Though current treatment is effective for localized high-grade osteosarcoma (HGOS) (70% 5 year overall survival,) the 20% of patients who present with metastatic disease remain difficult to cure, with overall survival of less than 20%.^2,3^.Progress has stalled in part because of the complex and heterogeneous alterations OS genomes typically harbor.^4^ While discrete subtypes can be distinguished from other pediatric cancers based on recurrent fusion proteins or cytogenetic abnormalities that guide therapy,^5^ osteosarcoma has largely eluded meaningful sub-characterization. Beyond a high frequency of *TP53* pathway alterations, an important characteristic of OS is the presence of complex genomic rearrangements (CGRs). To identify which mutational mechanisms contributed to the formation of CGRs, characteristic patterns of structural variants (SVs) and somatic copy number alterations (SCNAs) can be inferred from whole-genome sequencing (WGS) data. Recent work has identified an OS-specific mutational mechanism, termed Loss-Translocation-Amplification (LTA) chromothripsis, triggered by biallelic *TP53* inactivation through a single double-strand break, which in turn leads to a multi-generational genomic instability processes that results in oncogene amplification.^4^ The CGRs in osteosarcoma, which can contain hundreds of clustered SVs and segmental amplifications across multiple chromosomes, are established early in tumorigenesis and subsequently drive genomic instability and high levels of intratumoral heterogeneity, likely complicating treatment.^4^

Autochthonous, genetically engineered murine cancer models, which develop tumors in the same tissue as the original disease, have furthered our understanding of cancer development, evolution and response to therapy in the context of their native tumor microenvironment.^6–11^ However, whether murine cancer models recapitulate the genomic complexity observed in human tumors remains to be elucidated. Mouse models are often criticized as genomically simple as they usually have a lower tumor mutational burden (TMB) than their corresponding human cancers.^12–14^ This is exemplified by the *Trp53*^*fl/fl*^*Rb1*^*fl/fl*^ small cell lung cancer and KrasG12D-Trp53^fl/fl^ lung adenocarcinoma models, where the TMB in mice is 10-100 fold lower than humans.^14,15^ Several factors may contribute to the lower TMB detected in these mouse models: rapid development during weeks to months rather than years could limit accumulation of mutations;^16^ engineered, highly oncogenic initiating events like KrasG12D may minimize the fitness benefit of additional mutations;^14^ and early polyclonal diversification may generate many mutations below the limit of detection.^17^ Several groups have examined CGRs in autochthonous models but have found that CGRs are often rare (13% in Vk*Myc myeloma)^18^ or require engineered biallelic DNA repair defects rarely seen in the corresponding human cancers.^19^

To better understand the significance and therapeutic implications of CGRs in OS, we sought out an autochthonous murine model of OS that closely parallels the human disease.^20,21^ Prior studies of murine OS models reported complex karyotypic aberrations from spectral phenotyping.^20,21^ However, whether the same rearrangement mechanisms underpin the chromosomal aberrations detected in mOS models and human OS tumors remains elusive, thus limiting the relevance of mOS models for therapeutic development. To address this, we established a cohort of 234 mice based on a previously reported model of conditional *Trp53* and *Rb1* deletion in osteoblast progenitors, which we refer to as mOS (Fig. 1a).^20,21^ To assess the SV landscape of mOS, we performed high-depth, multi-region WGS of 35 matched primary and metastatic mOS tumors from 24 mice, and compared the genomic rearrangements detected in mOS with those found in human OS. Genomic analysis revealed frequent, often multichromosomal, CGRs with a level of complexity comparable to human OS that led to recurrent SCNAs in tumor suppressor genes (TSGs) and oncogenes, which we validated through transcriptional, proteomic and functional analyses.

**Figure 1.**
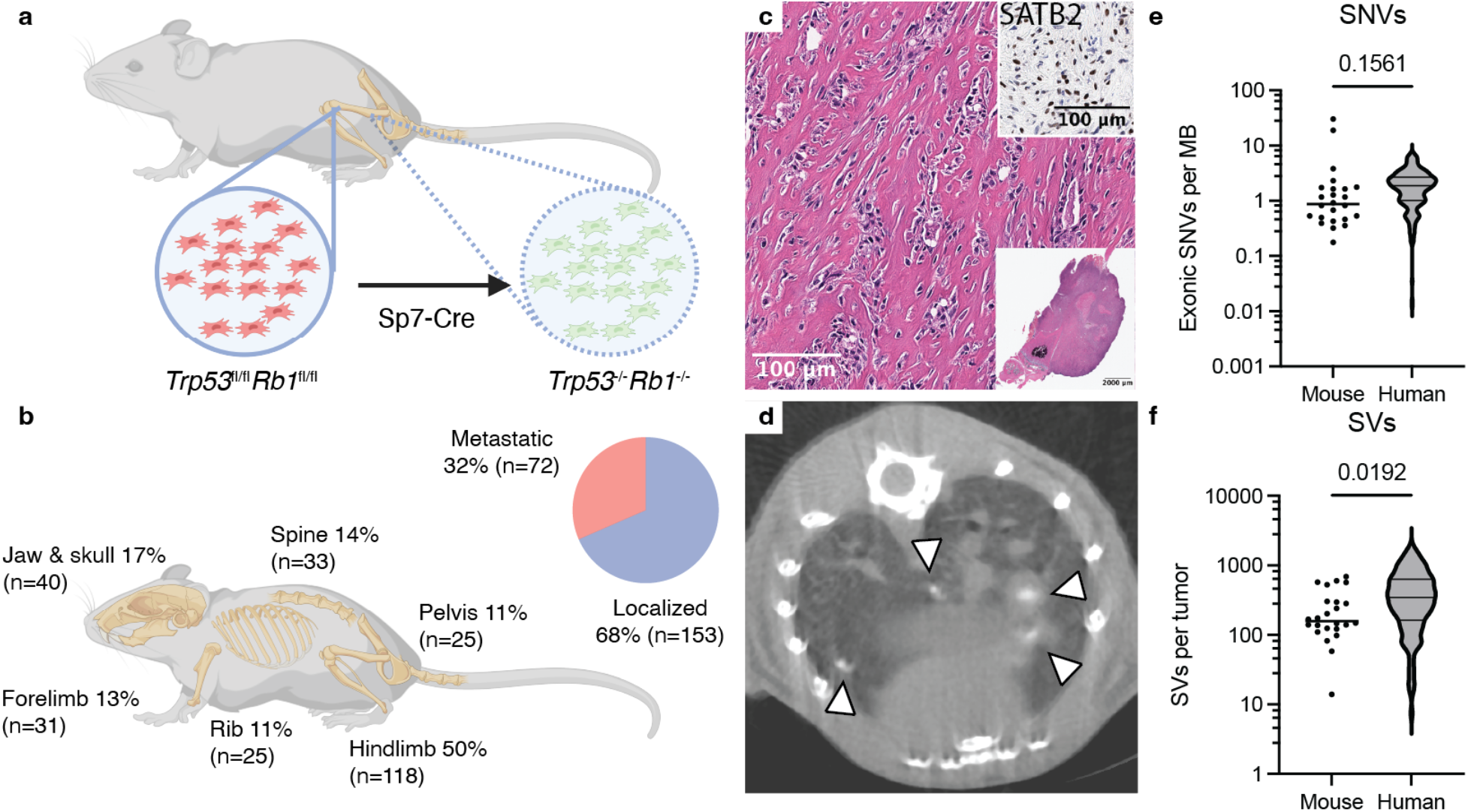
mOS models recapitulate clinical, pathologic and genomic features of human HGOS. **a)** Schematic of the genetically engineered murine osteosarcoma model used in this study. **b)** The distribution of physiological locations of tumors across our mOS cohort is shown (n=234, see **Table 1**). The total percentage exceeds 100% as 18% of mice had more than one tumor. **c)** Representative osteoblastic histology (H&E staining), with immunohistochemistry for SATB2 in the inset. **d)** Microcomputed tomography (µCT) of metastatic lung lesions. Arrowheads mark calcified metastatic sites. **d)** Somatic SNVs in coding exons per megabase (MB) from mOS tumors (n=24 unique cases), compared to all HGOS cases from Valle-Inclan et al.^2^ **e**) SVs per tumor (n=24), compared to all HGOS from Valle-Inclan et al.^4^ The p values in **e** and **f** were computed using the Welch’s t-test with a significance threshold set to 0.05

## Results

### mOS recapitulates histopathological and genomic features of human OS

To characterize the genomics of murine osteosarcoma and generate a model compatible with most common immunologic models, we generated a cohort of mOS on the C57Bl6/J background .All 234 of the mice in our cohort developed tumors that recapitulated key histopathological features of human osteosarcom: 1) Mice spontaneously developed firm, calcifying osteosarcomas; 2) the tumors were similarly distributed across anatomical locations as human HGOS with the majority of tumors (50%) developing in the hind limbs in the distal femur or proximal tibia (**Fig. 1b**); 3) histologically, tumors resembled human HGOS and produced osteoid and expressed the osteoblast transcription factor *SATB2* (**Fig. 1b**)^22^; and 4) approximately one third (32%) of mice developed detectable metastatic disease, most frequently in the lungs but rarely in the liver or kidney (**Fig. 1c**). Similar to prior reports,^20,21^ latency depended on genotype, as the median tumor onset ranged from 253 days (*Trp53*^*fl/f*^*Rb1*^*fl/fl*^) to 527 days (*Trp53*^*fl/+*^*Rb1*^*fl/+*^) (**Extended Data Fig. 1a**).

**Table 1.** Mouse OS cohort and set of tumor samples selected for sequencing.

|  | Overall mOS Cohort | Sequenced Cohort |
| --- | --- | --- |
| Tumor Samples | 359 | 35 |
| Primary | 287 (80%) | 27 (77%) |
| Lung Metastasis | 72 (20%) | 8 (23%) |
| Mice | 234 | 24 |
| Localized Disease | 153 (68%) | 10 (42%) |
| Metastatic Disease | 72 (32%) | 14 (58%) |
| Engineered Genotype |  |  |
| *Trp53<sup>-/-</sup>Rb1<sup>-/-</sup>* | 151 (65%) | 13 (54%) |
| *Trp53<sup>+/-</sup>Rb1<sup>+/-</sup>* | 12 (5%) | 1 (4%) |
| *Trp53<sup>-/-</sup>Rb1<sup>+/+</sup>* | 3 (1%) | 1 (4%) |
| *Trp53<sup>-/-</sup>Rb1<sup>+/-</sup>* | 58 (25%) | 8 (33%) |
| *Trp53<sup>+/-</sup>Rb1<sup>+/+</sup>* | 6 (3%) | 1 (4%) |
| *Other* | 4 (2%) | - |
| Median Tumor Onset | 245 days | 234 days |

The majority of mice (82%, 193/234) had a single primary bone lesion detectable at necropsy. This was surprising given the ubiquitous deletion of *Trp53* and *Rb1* in osteoblast precursors, which we confirmed by fluorescent reporter (**Extended Data Fig. 1b**), suggesting that successful osteosarcomagenesis requires additional, perhaps low probability alterations in the months following Cre-mediated deletion of *Trp53* and *Rb1*. To identify these mutations, we performed WGS of 35 tumor samples from 24 mice including both localized and metastatic sites from a range of engineered genotypes (**Table 1, Extended Data Fig. 1c**). Tumor samples were sequenced to high depth (average 110x) to assess CGRs and investigate the level of genomic heterogeneity in these tumors. In 20 mice (80%), a matched normal sample was sequenced to 30x coverage to filter out germline variants. For the remaining cases, we used a panel of normal samples to discount common germline variants as somatic (**Methods**). In aggregate, murine tumors had a TMB comparable to human HGOS^4^ (median 0.87 vs 1.87, *p*=0.16; Welch’s t test; **Fig. 1d**). There were significantly fewer SVs per tumor in mOS (median 158 vs 345, *p*=0.019, Welch’s t test; **Fig. 1e**). However, the range in number of SVs overlapped human HGOS (14 to 676, vs 7 to 1846), suggesting that despite the short latency relative to human disease,^4^ significant genomic complexity can develop in this model.

### Genomic landscape of mOS shows substantial inter-tumor variability and frequent CGRs

In-depth characterization of somatic SVs and SCNAs in mOS showed high inter-tumor variability and frequent CGRs (**Fig. 2**). All mOS tumor genomes harbored SCNAs, such as whole-chromosome losses and focal amplifications. Although the whole genome doubling (WGD) rate could not be inferred using WGS data from isogenic strains with few heterozygous SNPs, karyotypes and DNA content measurements from matched cell lines provided evidence for WGD events in all samples tested (3/3; **Extended Data Fig. 2**). This high rate was consistent with human HGOS where 99/141 (70%) of tumors had undergone WGD.^4^

**Figure 2.**
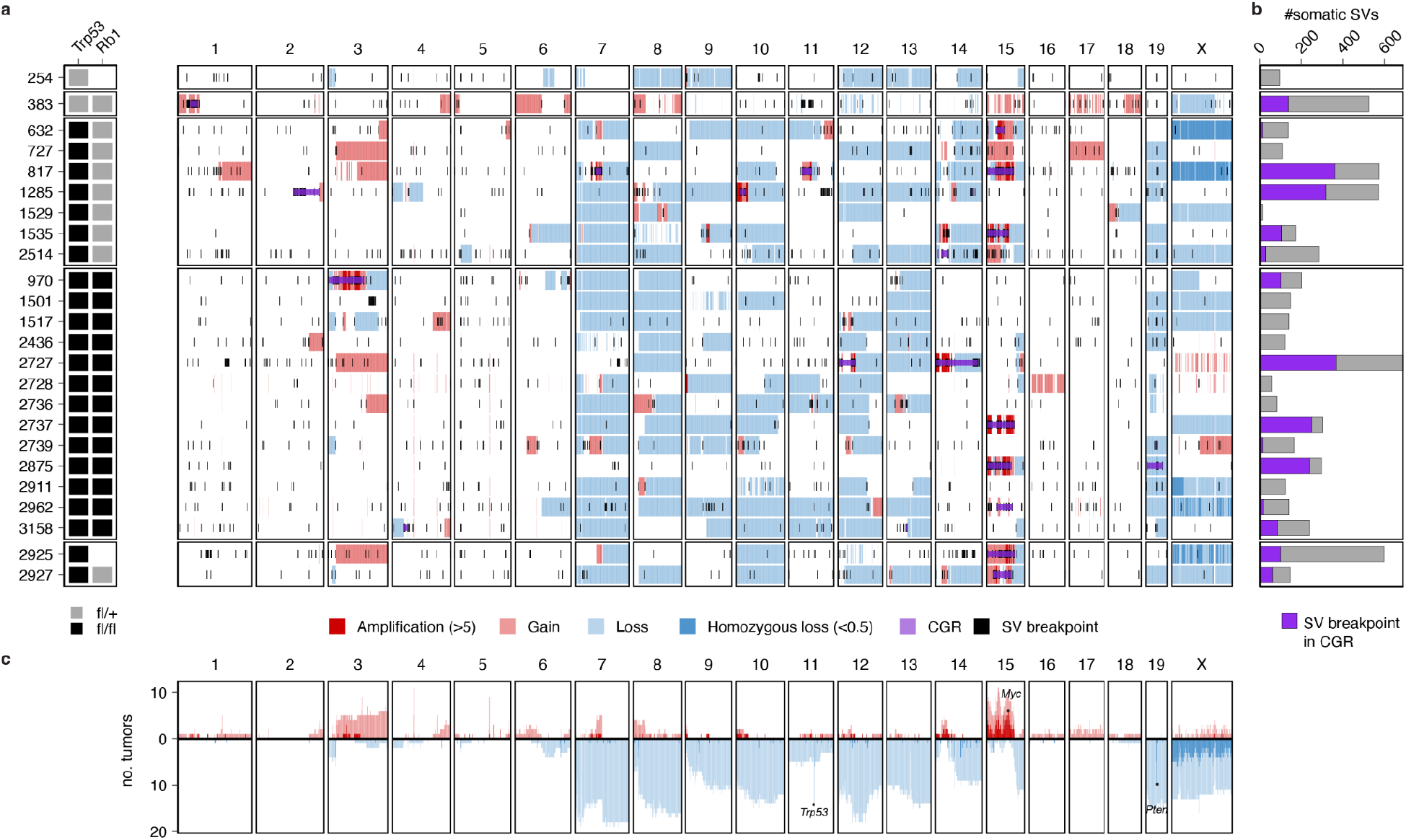
Structural variation landscape of mOS. **a)** Each row shows the SCNAs (gains in red, losses in blue) and SV breakpoints (black lines) detected in each tumor sample. CGRs are highlighted in purple. **b)** SV burden per tumor, with fraction of SV breakpoints mapping to CGR regions shown in purple. **c)** Number of tumors in the cohort showing amplifications (red) and losses (blue) across the genome.

In 15 of 24 (63%) primary mOS tumors, we identified clusters of interleaved SVs forming CGRs and substantially contributing to the total SV burden of the tumor, with a median 39% of SV breakpoints (range 5.4-82%) mapping to CGRs (**Fig. 2b, Supplementary Table 1**). In comparison, CGRs were detected in 131 of 141 (93%) human HGOS, including 45 of 46 (98%) cases with biallelic *TP53*/*RB1* inactivation.^4^ In mOS, the chromosome most affected by CGRs was chr15 (n=8/24 mice), which contains the *Myc* oncogene. Overall, CGRs were the main mutational mechanism through which segmental and focal amplifications were formed in this mouse model. In 8 cases (33%), mOS tumor genomes harbored CGRs involving multiple chromosomes, often with segmental amplifications, thereby generating complex karyotypes that bear resemblance to those observed in human OS (**Extended Data Fig. 3**). Overall, these results indicate that CGRs are a major driver of cancer genome evolution in both human OS and the mOS model analyzed in this study.

To investigate whether the CGRs formed in mOS tumors were generated by the same mutational mechanisms as the CGRs in human OS, we analyzed patterns of SVs and SCNAs in detail. In tumor 817, the co-occurrence of clusters of interleaved SVs, foldback inversions, segmental amplifications and terminal loss on chromosomes 7, 11 and 15 is consistent with a multigenerational rearrangement process involving both chromothripsis and breakage-fusion-bridge (BFB) cycles (**Fig. 3a-b**). A similar multi-chromosomal CGR was identified in tumor 1535 affecting chromosomes 9, 14 and 15 (**Fig. 3c-d**). In both mice, these CGRs underpin the amplification of oncogenes, including *Myc, Igf1r, Smad3* and *Duxbl1* (**Fig. 3a, d**). The SCNA and SV patterns of CGRs mOS show a clear resemblance to CGRs typically found in human OS arising from LTA chromothripsis (**Fig. 3e-h**). In human OS, it has recently been elucidated that LTA chromothripsis underpins oncogene amplification and the formation of the highly complex karyotypes observed in high-grade osteosarcomas.^4^ The CGRs we detected in mice 817 and 1535 harbor the features of LTA chromothripsis, suggesting they might arise from the same mutational mechanism (**Fig. 3, Extended Data Fig. 4**).

**Figure 3.**
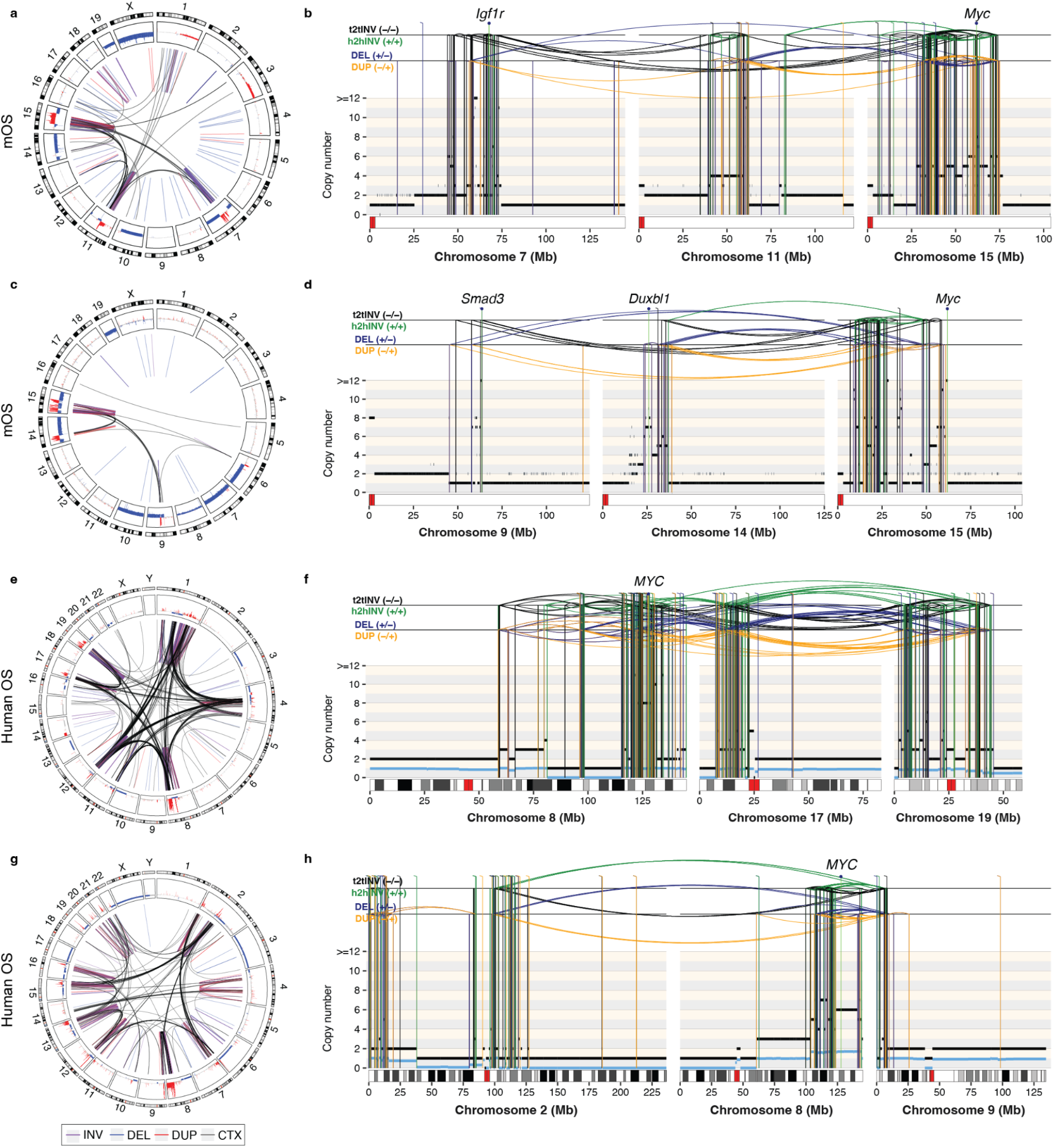
Representative examples of complex genomic rearrangements in mOS and human HGOS amplifying oncogenes. **a)** Circos plot depicting SVs and SCNAs detected in tumor 817. **b)** Multi-chromosomal rearrangement profile plot for 817 with SCNAs (horizontal lines) and SVs (vertical lines) showing oscillations, interleaved SVs, focal amplifications and inter-chromosomal breakpoints consistent with combined chromothripsis and BFB cycles. Copy number values are rounded and shown relative to a baseline of 2n. **c-d)** Circos and rearrangement profile plots for tumor 1535 showing a multi-chromosomal CGR involving chromosomes 9, 14 and 15. **e-f)** Circos and rearrangement profile plots for a representative human HGOS with LTA chromothripsis. **g-h)** Circos and rearrangement profile plot for a representative human OS tumor with biallelic *TP53*/*RB1* inactivation, harboring a *MYC* amplification generated by a multichromosomal CGR. Circos plot tracks in **a**,**c**,**e**,**f** depict (from outside to inside): (i) chromosome ideograms showing Giemsa banding data; (ii) SCNAs relative to the tumor baseline ploidy represented as log2 fold-change copy number values, with gains and losses shown in red and blue, respectively; (iii) SVs depicted by lines and colored by SV types: inversions (purple), deletions (blue), duplications (red) and translocations (black). Rearrangement profile plots display purity-adjusted rounded copy number values. Minor allele copy number values (i.e., the copy number values for the least amplified allele) are displayed in blue. SVs are depicted by vertical lines and coloured according to the SV type: DEL (deletion-like rearrangements) in blue, DUP (duplication-like) in orange, h2hINV (head-to-head inversion) in green, and t2tINV (tail-to-tail inversion) in black.

### Loss of *Trp53* and *Rb1* leads to CGRs in murine small cell lung cancer

Because deletion of *Trp53* and *Rb1* in the lung via adenoviral Cre recombinase delivery generates murine small cell lung cancer (mSCLC), we next examined whether these SCLC tumors developed similar complexity to mOS.^15^ To this aim, we reanalyzed WGS data from our group^15^ and others^13^ totalling 14 mSCLC tumors and found a higher number of SVs than in mOS (mean 545 vs 248, p=0.0184, Welch’s t test), potentially reflecting the longer latency of mSCLC (median 425 days) relative to mOS (median 253 days) (**Extended Data Fig. 5a**). Many (11/14, 79%) of the mSCLC tumors harbored interleaved SV clusters indicative of CGRs (**Extended Data Fig. 5b**). Interestingly, 12/14 (86%) tumors had segmental amplifications chromosome 4 involving the oncogenes *Mycl, Nfib* and *Trit1*, all of which have been associated with human SCLC (**Extended Data Fig. 5b-c)**.^**13**,**23**,**24**^ Overall, these results suggest that the concurrent loss of *Trp53* and *Rb1* in both mOS and mSCLC predisposes to CGRs that amplify oncogenes, and that the tissue context selects for specific amplifications, as recently suggested by forward genetic screens.^25^

### Multi-region sequencing shows that metastatic mOS evolves via punctuated evolution

Metastatic spread is the strongest predictor of poor outcome in human OS.^1^ However, research into how tumors confer metastatic potential has been limited by the difficulty of acquiring metastatic samples from human donors. Recent multi-region WGS of human OS revealed that LTA chromothripsis drives punctuated evolution and ongoing genomic instability.^4^ To assess the role of punctuated evolution in mOS tumor evolution, we characterized two metastatic mOS cases by multi-region WGS (**Fig. 4**). In each case, we classified SNVs and SVs based on whether they were detected in all (truncal), a subset (shared) or just one (private) of the tumor samples. Next, we constructed tumor phylogenies based on SNVs and SVs to analyze cancer genome evolution (**Methods**).

**Figure 4.**
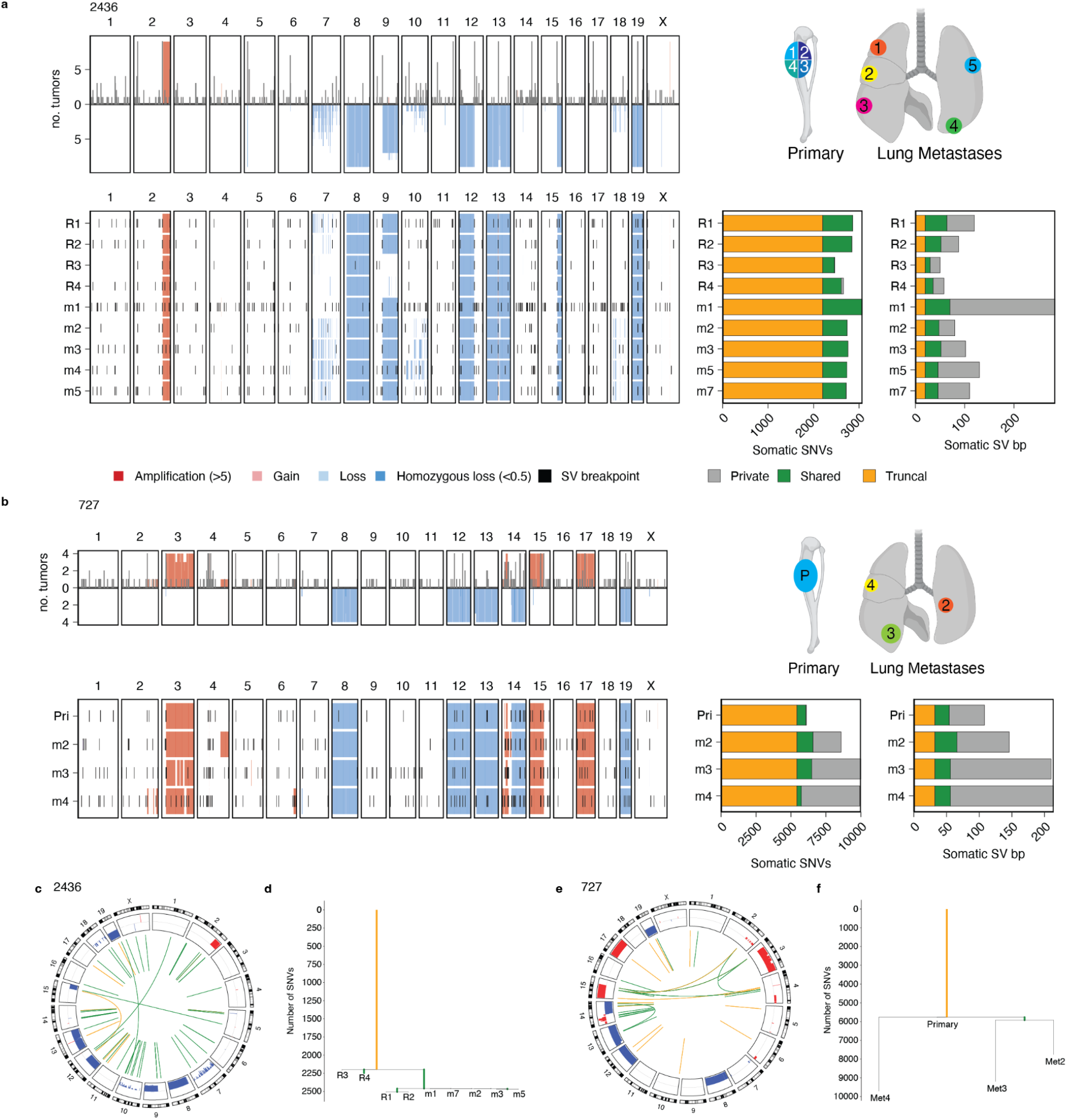
Punctuated evolution in mOS. **a-b)** Analysis of sequencing data from multiple tumor sites and regions from mice 2436 **(a)** and 727 **(b)**. Top panel with summary of recurrent SCNAs and SV breakpoints. Right diagram shows the approximate anatomical location of sampling sites. The bottom panel depicts the SCNAs and SVs detected in each sample. The bar plot on the right reports the number of private, shared and truncal SNVs and SVs. **c-d)** Circos plots representing shared alterations in samples from mice 2436 **(c)** and 727 **(d)**, depicting truncal (orange) and shared (green) SVs. Recurrent gains and losses are shown in red and blue, respectively, with a height proportional to the number of samples in which they occur. **e-f**) Phylogenetic trees based on SNVs for 2436 (**e**) and 727 (**f**). Met: metastatic sample; Pri: primary tumor; R: tumor region.

Mouse 2436 had a large primary tibia tumor, which we divided into 4 samples for WGS, and innumerable lung metastases, of which we sequenced the 5 largest (**Fig. 4a**). Large SCNAs were identified in chromosomes 2, 8, 9, 12, 13, 15 and 19. The genomic location and copy number of these SCNAs was highly consistent across all samples, except for a large deletion in chromosome 9 that was not identified in tumor region 3 (R3) or R4 (**Fig. 4a, c**). Overall, the average chromosomal copy number computed using 1Mb bins was highly concordant across samples (98% median pairwise cosine similarity, range 92-98%), and recurrence analysis also showed that large SCNAs were established early in tumor evolution. Phylogenetic analysis revealed that a median 80% of SNVs were truncal, indicating that the most recent common ancestor harbored the majority of SNVs. In this mouse, the phylogenetic analysis shows clustering of the metastatic samples with regions R1 and R2, while regions R3 and R4 diverged earlier (**Fig. 4d**). Although the samples acquired from this mouse harbored relatively few SV breakpoints (median 78 with allele fraction values of 0.1 or higher), we identified more private than truncal SVs (median 28 vs 20) which may suggest ongoing SV accumulation as observed in human HGOS (**Fig. 4a**). The SV burden of metastatic and primary samples was not significantly different (median 112 vs 73, p=0.18, Welch’s t test). Moreover, the short intermediate branches and long private branches of the SV phylogeny are in line with the rapid subclonal diversification described in human disease.^4^

The second multi-region case, mouse 727, included one primary sample from a tibia amputation and 3 lung metastases that were detected and collected 6.5 and 8 weeks after amputation, respectively (**Fig. 4b, Extended Data Fig. 6a-c**). Similar to 2436, the large chromosomal SCNAs were shared between the primary tumor and the lung metastases, as were more than half of the SNVs (59-92%, median 65%) and 19% of SVs, indicating that the substantial chromosomal aberrations were acquired in the most recent common ancestor. In total, we identified 34 truncal and 38 shared SVs, but the metastases in this mouse carried more private mutations (range: 36-96 SVs, **Fig. 4b, d**). The SNV and SV phylogenetic analyses were consistent with each other and showed that each metastasis arose from the primary tumor independently and not from metastasis-to-metastasis seeding.

Analysis of genomic regions with variable copy number patterns across tumor samples from mouse 727 revealed a CGR on chromosome 14 underpinning the amplification of oncogene *Duxbl1* (**Extended Data Fig. 6d-g**). We detected the rearrangements associated with this CGR in all samples. However, the number of copies of the amplifications generated by this CGR was low in the primary tumor compared to the metastases. Moreover, the SCNA patterns of Met2 and Met3 were most similar to each other and differed from Met4, corroborating the phylogenetic trees constructed from SNVs and SVs. Together, these results indicate that the genomic region affected by the CGR amplifying *Duxbl* underwent multi-generational genomic evolution, which is consistent with the instability generated by CGRs in human OS.^4^

Overall, our findings are in line with the patterns of punctuated evolution observed in human OS characterized by bursts of genomic instability that generate rapid clonal diversification, oncogene amplification and high levels of intra-tumour heterogeneity.^4^ Despite the engineered *Trp53* and *Rb1* knockout, the mOS tumors maintained the global karyotype of the most recent common ancestor, as defined by large-scale chromosomal gains and losses, whilst also accumulating SVs, thereby recapitulating the evolutionary dynamics of human osteosarcoma.

### Driver mutations in mOS resemble human HGOS

Next, we investigated whether the mutations acquired in the mOS model affect the same biological processes as in human HGOS. For a comprehensive assessment of genetic mutations, we considered somatic SNVs, indels, SV breakpoints and focal copy number changes in genes previously implicated in human HGOS (**Fig. 5a-b**). The most frequent somatic mutations in mOS were disruption of *Magi2* (n=14/24, 58%), *Naaladl2* (n=13/24, 54%) and *Pten* (n=12/22, 55% excluding engineered *Pten* cases 2925 and 2927), as well as amplification of *Myc* (n=6/24, 25%). In human HGOS, *MYC* amplification and *PTEN* loss are often discussed as putative driver genes, although their mutation frequency varies by study (e.g., 10-36% for *MYC* amplification and 6-17% for *PTEN* loss). ^4,26–28^ Most mOS cases (21/24, 88%) had at least one focal loss of a TSG or amplification of an oncogene implicated in human OS. In 7/8 mOS tumors without *Pten* disruption or *Myc* amplification, we identified amplifications and homozygous losses inside other cancer genes such as Il7r, Fbxw7, Yap1, Birc3 (**Fig. 5a, Supplementary Table 2**). Overall, from a panel of 947 genes reported as inactivated or amplified in prior studies of human OS, 775 were altered in at least one mOS case (82%), indicating generally high cross-species correlation.

**Figure 5.**
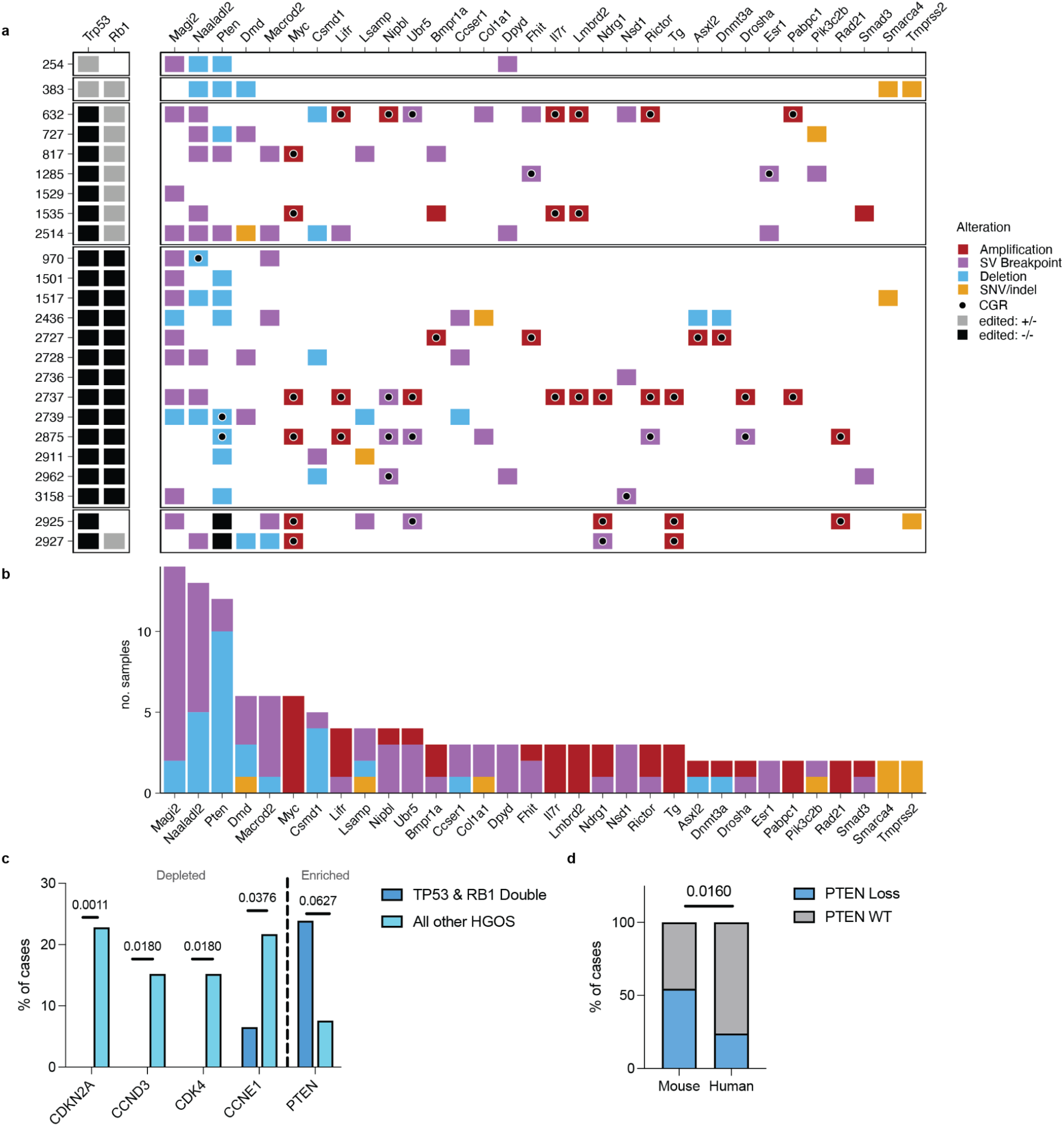
Genes mutated in human HGOS genes are frequently mutated in mOS. **a**) Panel of genes mutated in both HGOS and mOS sorted by frequency of mutation. The left column depicts engineered murine genotypes: one copy edited (+/-) in grey and both copies edited (-/-) in black. The gene level alterations are categorised as follows. Amplification: 5 or more copies; deletion: 0.5 or less copies and at least 0.5 copies below the chromosome average copy number; SV breakpoint: SVs mapping to exons and translocation breakpoints anywhere in the gene body; and SNVs and indels predicted to have a moderate or high impact as per Variant Effect Predictor. **b)** Gene level alterations are summarized as frequency in cohort. *Pten* frequency excludes edited cases 2925 and 2927 (**c**) Depletion and enrichment of gene mutations in cases with biallelic inactivation of *TP53* and *RB1* compared to all other HGOS cases. Rates were compared by Fisher’s exact test with Benjamini-Hochberg (BH) false discovery rate reported. **d**) Rate of *PTEN* loss in mouse and human HGOS cases with biallelic inactivation of *TP53* and *RB1*,^2^ compared by Fisher’s exact test.

However, some of the most frequently amplified and disrupted genes in human HGOS were not altered in mOS. Cell cycle regulators *CCNE1, CCND1 and CDKN2A* are frequently altered in human HGOS,^4,26,27^ but we did not detect any alterations of the murine orthologs in mOS. In mOS, the model’s engineered *Rb1* loss may reduce the need for and/or the evolutionary advantage of additional alterations dysregulating the cell cycle. To test the hypothesis that *Rb1* loss might mitigate the need for other mutations in cell cycle pathway genes, we compared our results to the mutation data from the human HGOS cohort (n=141) reported by Valle Inclan *et al*.^*4,26–28*^ Indeed, while these cell cycle alterations are common in human HGOS overall, in the subset of HGOS tumors with *TP53* and *RB1* mutations (n=46), they are rarely seen (**Fig. 5c, Supplementary Table 3**).^4^ In those patients, *PTEN* loss occurs significantly more often, while *CCND3* and *CDK4* amplification and *CDKN2A* loss are seen less frequently, paralleling the findings in mOS. Nevertheless, *Pten* loss in mOS (12/22, 55%) was significantly more common than *PTEN* loss in human *TP53/RB1* double mutant tumors (11/46, 45%, p=0.0125 Chi-square, **Fig. 5d**).

Next, we compared the prevalence of oncogene amplifications and their co-occurrence with CGRs in mOS and human HGOS.^4^ The prevalence of oncogene amplification was similar between mOS (11/24, 46%), human HGOS (79/141, 56%) and human HGOS with biallelic *TP53* and *RB1* inactivation (19/46, 41%). For all oncogenes in the gene set from Valle Inclan et al.^4^, we assessed whether their amplification coincided with a CGR on the same chromosome. The frequency of oncogene amplifications mediated by CGRs was high in mOS: 35 of 38 oncogene amplification events (92%) co-occurred with a CGR compared to 81% (152/188) and 60% (18/30), respectively for human HGOS and human HGOS with biallelic *TP53*/*RB1* inactivation. Thus, although the rate of CGRs in mOS is lower than in human HGOS (63% vs 93%, respectively), these findings emphasize the prominent role of CGRs in the formation of oncogene amplifications in osteosarcomagenesis across species.

### *Pten* is recurrently disrupted in mOS and contributes to tumorigenesis

Since WGS detected the inactivation of *Pten* by SCNAs or SVs in 59% of mOS tumors, we next sought to assess the contribution of PTEN inactivation to osteosarcomagenesis. Immunohistochemical (IHC) assessment of PTEN protein expression confirmed WGS results (**Fig. 6a, Extended Data Fig. 7 a-b**). All tumors with predicted biallelic loss and one tumor with biallelic SVs had no detectable PTEN protein relative to adjacent healthy tissue, while tumors with wild-type *Pten* status or heterozygous loss had preserved protein expression (**Fig. 6a, Extended Data Fig. 7 a-b**). We derived 4 tumor cell lines from mOS tumors, one of which was *Pten*-null by WGS and IHC (Line 383). Relative to cell lines heterozygous or wild type for *Pten*, Line 383 had constitutive phosphorylation of AKT Thr308, the expected result of unopposed PI3K activity, as well as downstream phosphorylation sites AKT Ser473 and pS6 Ser240/244 (**Fig. 6b-c, Extended Data Fig. 7c)**. By contrast, the other three cell lines had pAKT Thr308 and S473 detected near the level of the no-primary antibody staining control. pS6 Ser240/244 levels were increased over basal levels, possibly reflecting the contribution of other signaling pathways in those cell lines.^29,30^

**Figure 6.**
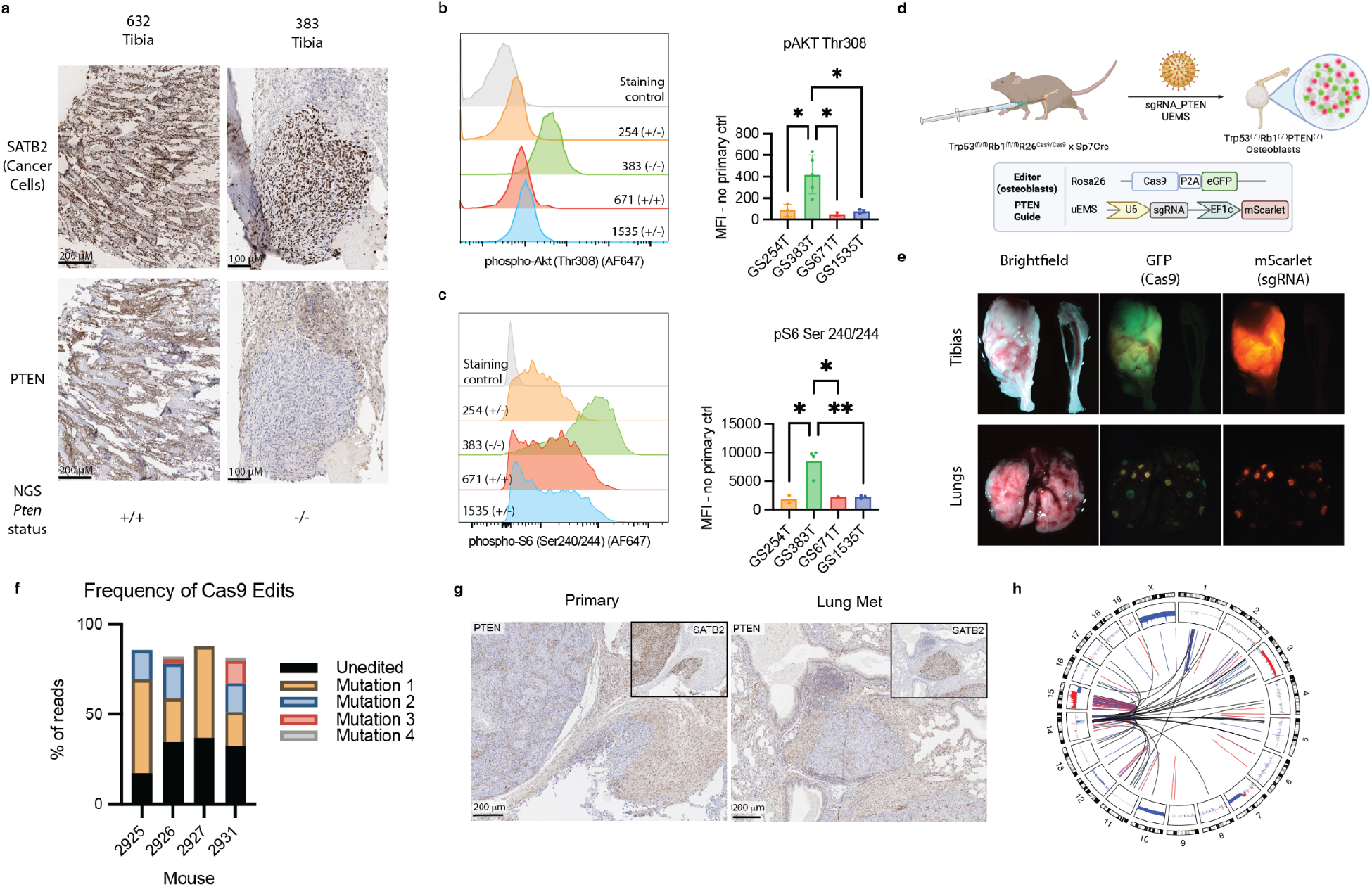
Recurrent *Pten* loss can drive mOS tumor formation. **a)** IHC validation of WGS-determined *Pten* status on formalin-fixed, paraffin embedded (FFPE) tumor samples. **b**) Phosphoflow cytometry of mOS-cell line basal phospho-AKT Th308 and **d)** phoshpho-S6 Ser240/244. **e)** Experimental design for tumor initiation by *Pten* guide using CRISPR/Cas9. e) Brightfield and fluorescent imaging of gross tumor primary and two metastatic sites. **f)** Frequency of all edited alleles >1% by WGS in each edited mouse. **g)** PTEN IHC on edited primary and metastatic sites from mouse 2925. **h)** Circos plot showing the genomic rearrangements underpinning the spontaneous amplification of Myc in mouse 2925. In **c** and **d**, one-way Anova (* p<=0.05, ** p<=0.01).

Despite universal deletion of *Trp53* and *Rb1* in all osteoblast lineage cells, 82% of mice had only one detectable tumor. Coupled with the long latency, this suggested successful osteosarcomagenesis was a rare event requiring additional lesions beyond the engineered deletion of *Trp53* and *Rb1*. We hypothesized that if *Pten* loss was contributing to tumor initiation, localized deletion in osteoblasts would allow spatio-temporal control of tumor formation in this otherwise spontaneous model. Thus, we next bred mOS mice to a conditional Cas9 allele and delivered *Pten*-targeting sgRNA to the left proximal tibia via lentiviral infection (**Fig. 6d**). Four of six mice developed left tibial osteosarcomas with distal metastases in the lung and, in one case, in the liver. All four of these tumors expressed the *Pten*-targeting guide (indicated by mScarlet positivity) and Cas9 (GFP staining) at both primary and metastatic sites (**Fig. 6e**). Timing of tumor formation was variable, ranging from 230 to 376 days old depending on *Rb1* status (**Supplementary Table 4**). *In vivo* editing was confirmed by amplicon sequencing (Crispresso2) and, in 2 cases, WGS. By all modalities, editing efficiency ranged from 56 to 64%, which corrected to 70-77% when accounting for cancer cell fraction in WGS samples (**Fig. 6f, Extended Data Fig. 8a, Supplementary Table 4**). These edits appeared to be selected for early in tumorigenesis, as each mouse had between 1 and 3 dominant alleles (**Fig. 6f, Extended Data Fig. 8a**). All tumors had complete loss of PTEN protein by IHC in primary and metastatic sites (**Fig. 6g, Extended Data Fig. 8b**). Of the remaining two mice, one died of non-cancer causes before any tumors developed and the other developed a jaw tumor without evidence of editing at 280 days of age. Interestingly, both cases analyzed by WGS had also developed CGRs on chromosome 15 resulting in *Myc* amplification (**Fig. 6h, Extended Data Fig 8c-d**). Though this cohort was too small to assess whether editing accelerates tumorigenesis, the spatial control and clonality of edits suggest that PTEN loss contributed to osteosarcomagenesis in these cases, consistent with the relatively high rate of *Pten* inactivation detected by WGS across the cohort.

### Complex genomic rearrangements amplify *Myc*

High oncogene amplification in ecDNA can result from chromothripsis and other CGRs.^31–33^ Across a diverse cohort of cancers, oncogene amplification in ecDNAs was associated with more aggressive disease, therapy resistance and worse overall survival than focal amplification.^32^ The presence of ecDNA has long been appreciated in human OS,^34^ and also observed in murine^35^ and canine OS.^36^ Because *Myc* amplification is associated with a poor prognosis in human OS,^37,38^ we were particularly interested in evaluating the development of *Myc* amplification in our mOS cohort. In six mice, we detected amplifications of *Myc* ranging from 5 to >100 copies (**Fig. 7a**). All of these amplifications were generated by CGRs mapping to chromosome 15, as exemplified by tumor 2875 (**Fig. 7b, Extended Data Fig. 3**). DNA fluorescence in-situ hybridization (FISH) on FFPE tissue confirmed *Myc* amplification (**Fig. 7c-d**). Remarkably, tumor 2875 with >100 *Myc* copies showed a very heterogeneous distribution of *Myc* puncta across cells, suggesting that *Myc* amplification may occur in extrachromosomal circular DNA (ecDNA). Based on the high copy number and high cell-to-cell variance, tumor 2875 was categorized as ecDNA positive by the scAMP model^39^ and the other amplified cases as ecDNA negative (**Extended Data Fig. 9b**). This categorization was further supported by inspection of individual nuclei in case 2875, in which *Myc* puncta in tumor cells were not associated with centromere 15 signals but were instead found in clusters throughout the nucleus (**Fig. 7e**).

**Figure 7.**
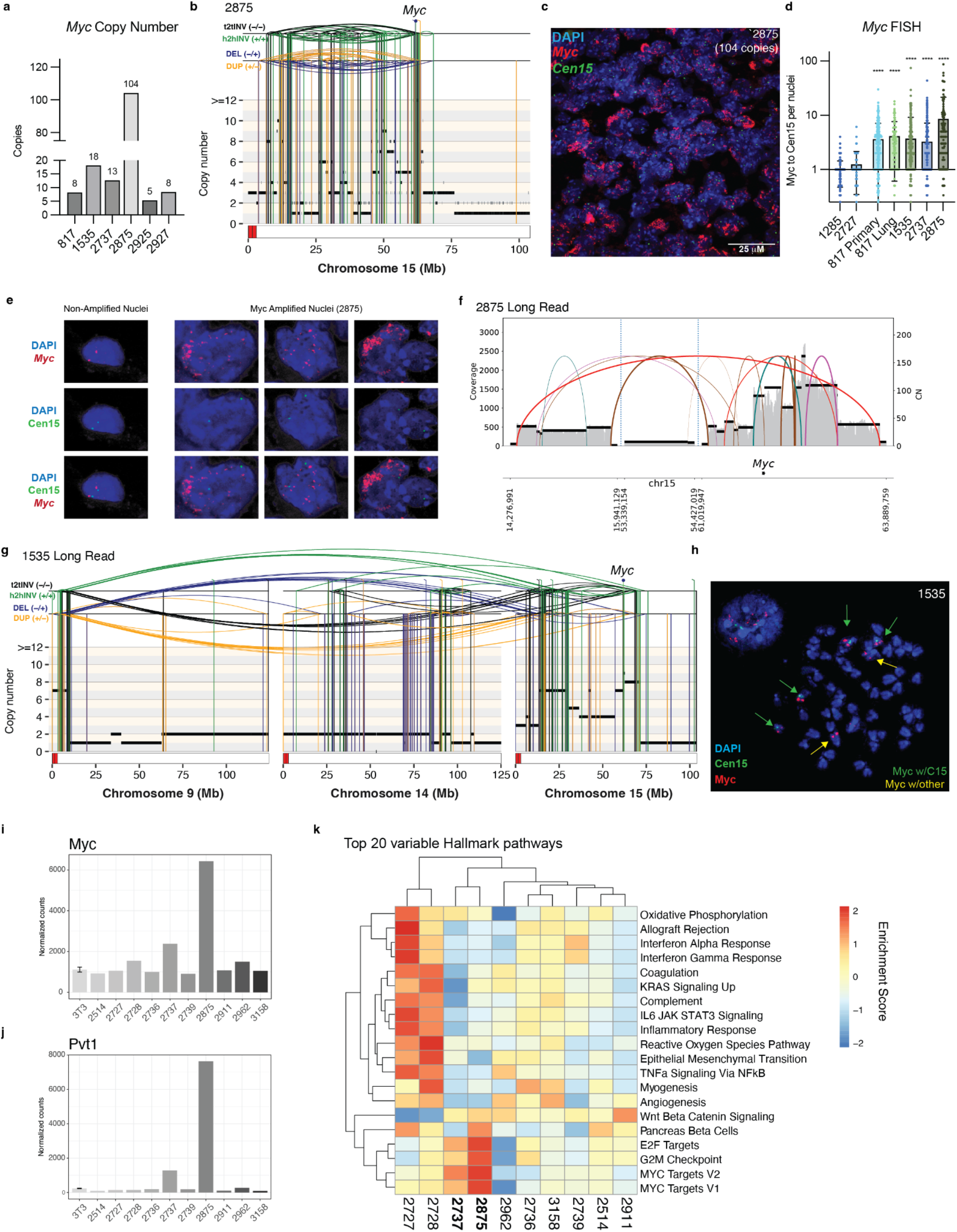
Recurrent *Myc* amplification in CGRs. **a)** *Myc* copy number in cases with amplification (>5). **b)** Representative rearrangement profile plot of tumor 2875. **c)** FISH on primary tumor sample from tumor 2875, quantified in **d)** by normalizing *Myc* puncta per centromere 15 puncta per nuclear area. **e)** Representative individual nuclei from tumor 2875. **f)** Breakpoint graph of 2875 amplicon from long-read sequencing. **g)** Rearrangement profile plot of 1535 derived from long-read sequencing. **h)** Representative metaphase FISH from 1535. **i-h)** *Myc* **(i)** and *Pvt1* **(j)** transcript from bulk RNA sequencing of 10 tumor samples and osteoblast reference line (MC3T3). **k)** Gene Set Variation Analysis of top 20 variable Hallmark Pathways among tumor bulk RNA-seq samples. Bolded samples are *Myc* amplified.

To corroborate the FISH findings, we attempted to reconstruct the ecDNA amplicons to disentangle putative ecDNA molecules from the CGRs that they potentially arose from. This prompted us to generate complementary PacBio long-read sequencing data for three tumors with varying degrees of *Myc* amplification: 2875, 1535 and 2737. Using CoRAL,^40^ which enables amplicon assembly from long-read sequencing data, we reconstructed multiple putative ecDNA structures in tumor 2875 (**Fig. 7f**). By contrast, ecDNA amplicons were not detected in 1535 or 2737. Instead, in the cell line derived from tumor 1535, long-read sequencing detected SVs connecting a centromeric amplification on chromosome 9 with two regions on chromosome 15, including *Myc* (**Fig. 7g**). This may represent neocentromere formation of a derivative chromosome, which is consistent with the multiple derivative chromosomes detected in karyotyping analysis that involve the chromosome band harboring the Myc gene (chr15(D2)) (**Extended Data Fig. 3d**). Indeed, metaphase DNA-FISH revealed intrachromosomal *Myc* amplification on chromosome 15 and an additional chromosome with *Myc* signal but no centromere 15 (**Fig. 7h, Extended Data Fig. 9c-d**), suggesting the presence of derivative chromosomes with intra-chromosomal amplification of *Myc*.

To understand the implications of CGRs amplifying *Myc* in the mOS model, we assessed the transcription in 10 murine OS tumors, including 2875 and 2737, by bulk RNA-seq. As predicted, the ecDNA+ tumor 2875 generated the highest number of *Myc* transcripts, though the ecDNA-tumor 2737 also had increased *Myc* transcripts compared to mOS without *Myc* amplification (**Fig. 7i**). *Pvt1*, which was predicted to be on multiple ecDNA cycles, was also strongly upregulated in 2875 and to a lesser degree in 2737 (**Fig. 7j**). Although *Pvt1-Myc* fusions can result in high levels of *Myc* expression in human cancers,^41^ we did not find any evidence of this fusion transcript in our data. The *Myc*-amplified cases did have increased *Myc* pathway activity based on Gene Set Variation Analysis (GSVA) of *Myc* and *E2F* target gene sets (**Fig. 7k**), underscoring the transcriptional significance of WGS-identified amplification. Collectively, evidence from sequencing data, FISH and karyotyping analyses suggests that *Myc* amplification in mOS is mediated by CGRs which can give rise to ecDNA as well as derivative chromosomes that increase the transcription of *Myc* and downstream target genes.

## Discussion

Despite poor outcomes, newly diagnosed patients with osteosarcoma today receive the same therapy they would have received forty years ago. Thus, new therapeutic approaches are desperately needed.^42^ Models that recapitulate human disease are valuable to address biological questions and test therapeutic strategies, especially given the scarcity and heterogeneity of patient samples with a rare disease like OS. Here, we described a cohort of mOS tumors that recapitulated the key clinical and histopathological features of human disease and performed deep genomic characterization. The mice spontaneously developed osteosarcoma after conditional deletion of *Trp53* and *Rb1*, which closely parallels human disease, where these disruptions are by far the most frequent recurrent alterations (78% and 48%, respectively) and are thought to be important drivers of tumor initiation.^4,26–28,43^ Furthermore, germline loss of *TP53* or *RB1* confers a 5-11% lifetime risk of developing OS.^44,45^ With biallelic, tissue-specific inactivation of *Trp53* and *Rb1*, all mice developed OS. Despite the global biallelic inactivation of *Trp53* and *Rb1*, most mice had only one primary tumor at necropsy nearly a year after *Trp53* and *Rb1* loss. The long latency prior to the development of a single tumor suggested that additional, low probability event(s) are required for osteosarcomagenesis.

Indeed, we found that all mOS tumors harbored additional somatic alterations in established oncogenes and TSGs. These alterations paralleled human HGOS cases with biallelic loss of both *TP53* and *RB1*. In mOS, *Pten* losses were common, while mutations in other cell cycle genes were not detected in line with human HGOS cases with biallelic loss of *TP53* and *RB1*. Mutations were also found in the most genes previously implicated in human HGOS, suggesting that this model recapitulates the wide range of genomic features of human HGOS. As expected for a model system, the parallel was not exact, likely due to species-specific biology. Mice have inherently long telomeres,^46^ and, consistently, we observed a low rate of mutations in genes involved in the alternative lengthening of telomeres pathway, the most common replicative immortality mechanisms in human OS.^4^ Specifically, we identified one case of *Top3a* amplification, two cases of homozygous *Atrx* loss, and no cases of *Daxx* loss.^47^ Similarly, differences in configurations and gene locations between mice and human chromosomes might affect the background rate of different mutational processes, especially those resulting in the formation of CGRs triggered by telomere crisis or micronucleation. Other recurrently altered genes in mOS were not mutated frequently or at all in human OS. One notable example was amplification of the *Duxbl* family of transcription factors on mouse chromosome 14, which has been reported in rhabdomyosarcoma in both mice and humans,^48^ and may represent novel OS biology for further study.

The functional significance of many of the gene level changes in OS has been inferred in past studies but has rarely been demonstrated genetically. Here, we developed a Cas9-based system to test the role of *Pten* loss, one of the most common alterations in our cohort. By locally deleting *Pten* in the left tibia, we were able to generate tumors in the injected bone with clonal edits at the predicted cut site. These tumors were aggressive, with 100% rate of lung metastases. The clonal selection suggested that *Pten* loss conferred a fitness advantage in osteosarcomagenesis and established its functional significance. Notably, at least half of these edited tumors also underwent CGRs on chromosome 15 amplifying *Myc*. This method can be extended to perform more systematic functional genomics in the development of this tumor type.

The genomic landscape of the murine OS model is one of the most complex seen across murine models of cancer, where CGRs, including chromothripsis, have been rarely reported.^18,19,49^ In murine OS, we most frequently observed CGRs on chromosome 15 resulting in *Myc* amplification through multiple mechanisms. In one tumor, DNA-FISH and long-read sequencing data revealed amplification through ecDNA, which has only been appreciated in a handful of spontaneous murine cancer models.^50,51^ In the other cases, the FISH *Myc* amplification patterns can be explained by a combination of derivative chromosomes and ecDNA, which is in line with the described mutational mechanisms of oncogene amplification in human HGOS^4^ and other cancers, such as breast cancer.^52^ Overall, the CGRs giving rise to oncogene amplifications in mOS mirrored those observed in human OS, suggesting that the same mutational mechanisms underpin the formation of CGRs in both species. Some of the multi-chromosomal CGRs resemble the recently reported LTA chromothripsis pattern, which is remarkable given that the acrocentric mouse chromosomes could be thought of as less prone to formation of dicentric derivative chromosomes and subsequent chromatin bridge formation. In mOS, it appears that multi-chromosomal genomic instability and/or the resulting multi-chromosomal rearrangements confer a significant fitness advantage.^46^ Though some cases were highly complex, numerically, mOS tumors still had fewer SVs than human OS. Since the multi-region sequencing data suggest an early cataclysmic genomic event followed by ongoing SV accumulation, the overall SV disparity may reflect the weeks to months timespan of murine tumors compared to the decade-long process of human tumor evolution, variable background SV rates, smaller genome size, and variable sensitivity for SV detection in human and murine OS.

Human osteosarcoma has one of the highest rates of chromothripsis of any cancer type.^33^ As we have shown here in mouse and has recently been described for canine osteosarcoma,^53^ chromothripsis appears to be a cross-species hallmark of OS. Whether this predisposition is a function of the loss of *TP53* and *RB1*, the cell of origin or unique features of the bone microenvironment, remains to be determined. We reanalyzed WGS data from the murine SCLC model, which also requires deletion of *Trp53* and *Rb1*, and found that these tumors also developed CGRs. Unlike mOS, where CGRs preferentially involved chromosome 15, nearly all mSCLC cases involved chromosome 4, which harbors *Mycl* and *Nfib*, both key oncogenes in SCLC.^13,24^ The high rate of CGRs affecting oncogenes in both mOS and mSCLC suggests that loss of *Trp53* and *Rb1* predisposes to CGRs that are then subject to tissue-specific selection. Since tumors in both models have a long latency period before tumorigenesis, they present a unique opportunity to study precursor lesions harboring CGRs, including ecDNA and unstable derivative chromosomes. For example, single-cell WGS of bone cells before tumor formation could be used to establish the background rate of CGRs *in vivo* across different cell types or whether CGRs develop long before or immediately preceding tumorigenesis, which would further our understanding of tumor initiation and progression in genomically complex tumors.

Overall, we present a murine osteosarcoma model that recapitulates key features of human OS, including its genomic landscape riddled with CGRs that may contribute to tumor initiation and immune evasion. This autochthonous model and the transplantable, genomically characterized cell lines derived from it, can be used in future studies of osteosarcoma initiation and tumor-immune interactions. The model’s genetic tractability, exemplified by the *in vivo PTEN* editing we performed, makes it particularly suited to address mechanistic questions in the native tissue microenvironment. Given the shared cross-species genomic alterations, studies should integrate data from human samples with mechanistic and preclinical murine and canine experiments to develop new therapeutic approaches for this devastating disease.

## Acknowledgements

This work was supported by Ludwig Cancer Research (T.J.) and St. Baldrick’s Foundation (G.A.S.). G.A.S. was supported by the Damon-Runyon/St. Jude Pediatric Cancer Fellowship (G.A.S.). I.A.E.M.v.B. and I.C.C. thank EMBL-EBI for funding. We appreciate the support of the Swanson Biotechnology center FACS, Histology, Preclinical Modeling and Imaging cores, which are supported by NCI 5P30CA014051. We are grateful to members of the Jacks and Cortes-Ciriano labs for their helpful feedback throughout.

Provision of patients’ samples from the RNOH was made possible through the Royal National Orthopaedic Hospital Pathology Department and the Research and Development Department, The Rosetrees Trust, Skeletal Cancer Trust, Sarcoma UK, The Bone Cancer Research Trust, and the Pathological Society of Great Britain and Ireland, over the last two decades. The project was also supported by the National Institute for Health Research, UCLH Biomedical Research Centre, and the UCL Experimental Cancer Centre. We thank the patients and their families for their participation in this study. This research was made possible through access to data in the National Genomic Research Library, which is managed by Genomics England Limited (a wholly owned company of the Department of Health and Social Care).^54^ The National Genomic Research Library holds data provided by patients and collected by the NHS as part of their care and data collected as part of their participation in research. The National Genomic Research Library is funded by the National Institute for Health Research and NHS England.

## Data and code availability

The mouse sequencing data generated in this study will be available in the Sequence Read Archive (SRA) repository upon publication.

WGS data from human osteosarcoma patients enrolled in the 100,000 Genomes Project can be accessed via Genomics England Limited following the procedure described at https://www.genomicsengland.co.uk/about-gecip/joining-research-community/. Applicants from registered institutions can apply to join one of the Genomics England Research Networks. Dual consented genomic data (somatic and germline) were shared to RNOH and EMBL-EBI. Data linked to local clinical data (no NHS Digital or NHS England data) were shared by Genomics England with RNOH and EMBL-EBI. No analysis was undertaken in the Genomics England Research Environment or National Genomic Research Library.

Raw WGS data from the Gabriella Miller Kids First Pediatric Research Program: An Integrated Clinical and Genomic Analysis of Treatment Failure in Pediatric Osteosarcoma (project number 1 X01 HL 132378-01) are available at dbGaP under accession code dbGaP: phs001714.v1.p1. The WGS data from the Therapeutically Applicable Research to Generate Effective Treatments (TARGET) program is available at dbGaP under accession code dbGaP: phs000218. The raw WGS data from the Behjati et al.^4,26–28,43^ and MD Anderson Cancer Center^55^ cohorts are available at the European Genome-Phenome Archive (EGA) under accession numbers EGAD00001000147 and EGAS00001003247, respectively. PCAWG analysis results are available at https://dcc.icgc.org/releases/PCAWG. The raw WGS data generated by TCGA are available at dbGaP under accession code dbGaP: phs000178

The code used to process the data, generate and visualize the results presented in this manuscript will be made available on github upon publication.

## Methods

### Mice

All animal studies described in this study were approved by the MIT Institutional Animal Care and Use Committee. mOS mice were rederived by backcrossing the previously developed Rb^fl^ mouse (B6;129-Rb1^tm3Tyj/J^ Jackson #008186) to C57Bl6/J (Jackson #000664) and subsequently breeding to Trp53fl mice (B6.129P2-Trp53tm1Brn/J, Jackson #) and Sp7 Cre Mice (B6.Cg-Tg(Sp7-tTA,tetO-EGFP/cre1Amc/J, Jackson #006361). The resulting mOS mouse was subsequently crossed to the Rosa26^lslCas9^ mice (B6J.129(B6N)-Gt(ROSA)26Sor^tm1(CAG-cas9*,- EGFP)Fezh/J^ Jackson #026175) to generate mOS-Cas9 animals. Animals were monitored weekly for tumor formation. C57Bl6/J (Jackson #000664) and Rag2 deficient (Rag2^tm1Fwa^ Taconic #601) mice were used as orthotopic transplant recipients. Rosa26-lsl-TdTomato mice (B6.Cg-Gt(ROSA)26Sort^m14(CAG-tdTomato)Hze^/J, Jackson #007914) were bred to Sp7-Cre to assess Cre-labeled tissues.

### Whole genome sequencing

Tumors, tumor regions or individual metastases were flash frozen in liquid nitrogen and manually pulverized (BioSpec BioPulverizer 59012N). The tail tip was digested with proteinase K for healthy control DNA. Genomic DNA and RNA were isolated using an AllPrep DNA/RNA Mini Kit (Qiagen #80204) or for DNA only, DNAeasy Blood and Tissue Kit (Qiagen #69504) as per manufacturer’s instructions. Briefly, genomic DNA (0.5 μg) was used to prepare sequencing libraries with the TruSeq DNA PCR-Free High Throughput Library Prep Kit (cat. #20015963, Illumina) as per the manufacturer’s instructions, with the exception of 2 samples with low DNA quantities (2436_met5, 2436_met7). In those cases, the TruSeq DNA Nano High Throughput Library Prep Kit (cat. #20015965, Illumina) was used. Tumor samples were sequenced to 100x coverage and healthy controls to 30x on a NovaseqX (150 bp paired-end reads). Library preparation and sequencing were performed at Psomagen (Rockville, MD).

Sequencing reads were mapped to the reference mouse genome version mm39 using BWA-MEM,^56^ after which Picard was used to mark duplicates and obtain quality metrics (**Supplementary Table 1**)

### Tumor purity estimation

To estimate the fraction of cells with edited *Trp53*, the coverage depth of the *Trp53* edit site (chr11:69474029-69481865) was compared to a broader region encompassing *Trp53* (chr11:69400000-69500000). The mean coverage depth of both regions was computed using the function coverage from the SAMtools package,^57^ restricting the analysis to properly paired reads (SAM flag 0×2) with a mapping quality ≥ 20 and base quality ≥ 20 (**Supplementary Table 1**). The obtained purity estimates were verified through manual inspection of sequencing reads.

### Detection of somatic copy number alterations

Somatic copy number alterations were inferred using CNVkit^58^ with default parameter values. When available, the matched normal sample was used. Alternatively, a panel of normals (PoN) constructed from all available control mouse samples (n=17) was used. Next, we removed segments that overlapped >=20% with low mappability or high signal regions as defined in the ENCODE blacklist^59^ in order to prevent spurious calls.

To facilitate downstream analyses, we inferred absolute copy numbers from the relative log2-fold copy number change provided by CNVkit using the following equation:

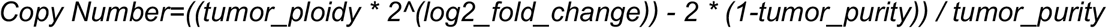

For cohort-wide analyses, we assumed a diploid tumor genome. Therefore, in tumors undergoing whole genome doubling followed by single-copy gains or losses, the estimated copy number values would be ∼2.5 and ∼1.5 copies, respectively. Given that a substantial fraction of tumors in our cohort has likely undergone whole-genome doubling, we used more lenient thresholds for the identification of single-copy gains (>=2.5 copies) and losses (<=1.5 copies). Yet, stringent thresholds were used to call amplifications (>=5 copies) and homozygous losses (<=0.5 copies).

Recurrent gains and losses were identified by binning the genome into 1 megabase (Mb) bins and counting the number of tumors with a gain or loss, respectively, in each bin. For mice 727 and 2436 we only included one sample of the primary tumor in the cohort-level analysis. Furthermore, to assess whether the overall SCNA profiles were similar across samples, we calculated the pairwise cosine similarity of average copy number across all 1 Mb bins across the genome.

### Detection of somatic structural variants

For the tumor samples with a matched control sample, structural variants (SVs) and indels were identified with SVABA^60^ using default settings. The variant allele fraction (VAF) was calculated by dividing the number of variant-supporting reads by the depth of coverage (AD/DP), as per developer’s recommendations.

For the tumor samples without a matched control sample, we intersected somatic SVs obtained from paired analysis with each of the healthy controls as follows: For each tumor-only sample (n=5), we inferred SVs using each of the healthy controls as background (n=17, excluding the *Pten* edited mice). Next, we selected the SVs that were identified in all of these 17 analyses as ‘somatic’ and never as ‘germline’. When comparing SVs between analyses, we considered two SVs as the same if their SV type (i.e., deletion, duplication or inversion) matched and the distance between the breakpoint coordinates was less than 10 base pairs. To detect somatic indels in tumor-only samples, we required the exact indel to be identified in all 17 tumor-normal comparisons.

CGRs were identified as clusters of at least 4 interleaved SVs (corresponding to 8 breakpoints) using ShatterSeek v1.1.^33^ The leftmost and rightmost breakpoints of the cluster were used to define the region of the CGR. To identify multi-chromosomal CGRs, we searched for inter-chromosomal breakpoints connecting interleaved clusters on different chromosomes. In addition, we considered two chromosomes to be involved in the same CGR if they were connected by more translocations than expected given the background rate of translocations in each tumor. (Fisher’s exact test < 0.01 after multiple testing corrections with Benjamini–Hochberg (BH)). Finally, all CGRs were verified via manual inspection.

### Detection of somatic single nucleotide variants and indels

For the tumor samples with a matched control sample, somatic short mutations including single nucleotide variants (SNVs) and indels were identified using Mutect2^61^ and MuSE.v1^62^ with default parameters. To filter out germline polymorphisms, we created a PoN using the data from all normal samples (n=17, excluding *Pten* edited mice). We considered as high confidence those SNVs detected by both algorithms. For tumor samples without a matched germline sample, we only ran Mutect2 and only considered as somatic those SNVs with a VAF > 0.1 after removing all SNVs detected in at least one germline control sample.

Indels were detected using Mutect2 and SVABA. We only considered as somatic those indels detected by both Mutect2 and SVABA and not detected in any of the germline controls.

### Tumor mutation burden analysis

For each tumor, the SV burden was calculated as the sum of SV breakpoints in all autosomes and chromosome X. To obtain the fraction of SV breakpoints in CGRs, the number of breakpoints overlapping the CGR regions were summed and divided by the total number of breakpoints in the tumor genome. The SNV burden was calculated using non-synonymous SNVs mapping to coding regions.

### Gene level alterations

For comprehensive analysis of gene level alterations, we combined SNVs, indels, SVs, amplifications and focal losses. To focus on the variants that are likely to affect the protein function, we annotated SNVs and indels using Variant Effect Predictor (VEP)^63^ and selected variants that were predicted to have a ‘moderate’ or ‘high’ impact and any of the following consequences: deleterious missense_variant, stop_gained, frameshift, start_lost, stop_lost. To identify SVs altering gene function, we considered all translocations and intrachromosomal SVs with at least one breakpoint mapping to an exon. To identify amplifications, we selected genes with a weighted average of 5 copies or more Similarly, to identify genes affected by focal deletions, we required a weighted average of 0.5 copies or less, in addition to at least 0.5 copies lower than the chromosome average copy number.

To establish a genes of interest panel for osteosarcoma, we combined gene lists from Valle-Inclan *et al*.^*4*^ and Sayles et al,^26^ and used the Homologene package to match the mouse gene names to human orthologs.^64^

To assess whether gene amplifications were potentially mediated by CGRs, we flagged amplifications if a CGR occurs in the same chromosome. This parallels the analysis of human tumors, where it is common to flag variants that occur in the same chromosome arm as a CGR, but since mouse chromosomes are acentric we used the full chromosome instead. In addition, a more specific annotation is provided in Supplementary Table 2 for amplifications that are directly overlapping with a CGR region.

### Phylogenetic analyses of multiregion data

To analyze the relationship between the multiple samples obtained from different tumor regions and metastatic sites of mouse 727 and 2436, we constructed phylogenetic trees from somatic SNVs with maximum parsimony using the parsimony ratchet method from the phangorn package.^65^ As input for the tree, for each mouse, we recorded in tabular format the presence or absence of each SNV in each sample. For this analysis, we considered high-confidence SNVs detected by both Mutect2 and Muse in at least one of the samples and a sequencing depth of at least 40x in all samples. In addition, we made a mutation call if we found at least 3 sequencing reads supporting the alternate allele with a mapping quality of at least 30 and a base quality of at least 20. We set the tree branch lengths to the number of truncal, shared and private mutations detected in each relevant subset of samples. The topology of the phylogenetic trees obtained was validated by bootstrapping analysis through subsampling of the mutations, and by manual inspection of sequencing reads across all samples at the loci to which mutations detected in a subset of samples were detected. In addition, we constructed phylogenetic trees from somatic SVs with an allele fraction >0.1 in at least one sample, using the Unweighted Pair Group Method with Arithmetic Mean method from the phangorn package.^65^ The SV phylogenies were consistent with the SNV-based phylogenies.

### Visualization

Figures were generated using R packages: ggplot2, circlize and ReconPlot.^66–68^

### Long read sequencing

DNA was extracted from frozen cell pellets (1535) or flash frozen pulverized tumor samples using the Nanobind Pan DNA Kit (PacBio 103-260-000) according to the manufacturer’s instructions. Libraries were prepared using the SMRTbell Prep Kit 3.0 (PacBio 102-182-700) as per the manufacturer’s instructions including short read elimination step. Circular Consensus Sequencing was performed on the Revio system (PacBio) with one sample per SMRT Cell. DNA extraction, library preparation and sequencing were performed at Psomagen (Rockville, MD). Sequencing reads were mapped to the reference mouse genome version mm39 using minimap2.^69^ For downstream analysis, split reads were extracted using samtools^57^ and data was visualized with IGV.^70^

Structural variants were inferred using SAVANA^71^ and SCNAs using CNVkit^58^ using default parameters. Amplicons, including ecDNAs, were reconstructed using CoRAL with default parameters after re-mapping reads to reference genome version mm10.^40^

### Bulk RNA sequencing and analysis

Tumor RNA was isolated in parallel with DNA as described above. RNA from cell lines, including MC3T3-E4 controls, was isolated with an RNAeasy kit. Total mRNA sequencing library was prepared by poly-A capture and sequenced by Novogene (Sacramento, CA) on a NovoseqX with ∼50 million reads per sample. Sequencing data were processed with nf-core/RNAseq v3.19.0 reference to GRCm38 (mm10) and analyzed with DESeq2. To identify RNA fusion transcripts, alignments to mm10 including chimeric transcripts were generated with STAR (v2.7.11b)^72^ and, together with the SVs identified above, analyzed with Arriba (v2.5.1)^73^ to identify fusion transcripts.

### Tumor cell line derivation and orthotopic transplants

A portion of viable tissue was physically dissociated by crushing through a garlic press (Amazon #B092D1DP4D) and then enzymatically digested per manufacturer’s protocol (Miltenyi Biotech 130-096-730). The resulting suspension was filtered through a 70 uM, centrifuged and resuspended in DMEM with 10% FBS and penicillin/streptomycin/glutamine and plated. Media was changed after 48 hours and then as indicated until plate became confluent (2-4 weeks). Cultures with sustained growth were expanded and orthotopically implanted into the tibia of immunocompetent and Rag2-deficient mice. Briefly, a pilot hole was drilled by inserting a 27 g, ¼” needle (BD) through the tibial plateau into the Mice were monitored for tumor growth by µCT and euthanized 8 weeks after implantation (or sooner if tumors exceeded 1 cm in size). Osteoid formation was confirmed on H&E staining. Cell Line 383T grew in immunocompetent with 100% penetrance, while 254T and 1535T only grew consistently in immunodeficient mice.

### Histology and immunohistochemistry

Tumor tissue was fixed in 4% paraformaldehyde overnight and decalcified in 0.5 M EDTA for 72 hours before paraffin embedding. For immunohistochemistry, slides were subjected to citrate based antigen retrieval prior to staining with the indicated antibodies (Supplementary Table 5) overnight at 4 C and detecting with the appropriate HRP-secondary.

### Karyotype & DNA content determinations

Cytogenetic analysis of cell lines was performed on 10 Giemsa banded metaphase cells from each line (Cell Line Genetics, Madison, WI). Cell line DNA content was determined by FACS analysis of PI staining relative to healthy murine splenocytes. Briefly, cells were fixed and permeabilized in ice cold 70% ethanol for 30 minutes, rehydrated with PBS, treated with RNAse A (Qiagen) for 10 minutes and mixed with propidium iodide (Invitrogen) before analyzing on a FACS Symphony A1.

### FACS signaling analysis

Tumor cell lines cultured in complete media were fixed and permeabilized in 2% PFA for 10 minutes, washed with FACS buffer and permeabilized by gradual addition of ice-cold 90% methanol and incubated at -20 C overnight. Samples were warmed to room temperature, rehydrated with FACS buffer and stained with primary antibody (Supplementary Table 5) for 30 minutes, washed, stained with fluorescent secondary antibody for 30 minutes and washed prior to acquiring on a BD LSR-Fortessa. Analysis was performed in FlowJo (BD, Version 11).

### In vivo Cas9-editing

Lentivirus (10 uL, 2.6e6 IU) encoding PTEN sgRNA (TCACCTGGATTACAGACCCG) and mScarlet reporter was injected into the left tibia intramedullary space by first passing a 25 g needle and then inserting a 31g Hamilton syringe into mOS-Cas9 mice 4-7 weeks old. Mice were monitored weekly for clinical signs of tumor and potential masses were assessed by uCT imaging. When tumors reached 1 cm in size, necropsy was performed to identify additional tumors and metastasis. Before further processing, tumors were imaged with a Nikon SMZ1500 stereomicroscope equipped with an X Cite series 120 fluorescence illumination system. Tumors were divided for genomic DNA extraction and IHC (both as described above). Metastatic sites were processed for IHC. Editing was assessed by Sanger Sequencing of the edited region (fwd CCGATACCCCTTACTGCCTCTG, rev gggacagcctggtctacaaagt) using TIDE^74^ and Amplicon sequencing (fwd TCCCTACACGACGCTCTTCCGATCTATTCCCAGTCAGAGGCGCTATG, rev GTTCAGACGTGTGCTCTTCCGATCTgggacagcctggtctacaaagt) using Crispresso2.^75^

### FISH and ecDNA prediction

FFPE sections (4 µm) were deparaffinized and subjected to antigen retrieval in citrate (pH 6.8) buffer at 95C for 30 minutes. Samples were digested with pepsin (1.5 mg/mL, Thermo Scientific) for 30 minutes at 37 C in 0.01N HCl. Autofluorescence was quenched with sudan black. *Myc* (Empire Genomics RP23-307D14 Orange) and *Cen15* (Empire Genomics RP23-333G9 Green) were pre-heated to 63 C for 5 minutes and then codenatured with the slide for 7 minutes at 75 C before incubating at 37 degree overnight. Samples were washed with 0.4x SSC/0.3% Igepal (Sigma Aldrich) for 2 minutes at 72 C and 2x SSC / 0.1% Igepal for 5 minutes before DAPI staining. Slides were imaged on an Evident FV4000 at 63x or 100x. Images were quantified with QuPath by DAPI-based nuclear region detection and subcellular spot detection to calculate the ratio of Myc to Cen15 probes per nuclear region. Nuclei with 0 Myc or 0 Cen15 puncta were excluded. The per cell output of *Myc*:Cen15 were analyzed with *scAMP* to infer ecDNA status using the following parameters: min_copy_number=2, max_percentile=99, filter_copy_number=0.^39^

### Reanalysis of Human OS Cases from Valle Inclan et al.^4^

*TP53* and *RB1* double mutant cases were identified from Supplementary Table 3. The 46 identified double mutant cases were manually confirmed to have biallelic loss of both genes in the original spreadsheet. Incidence between groups was compared by Fisher’s exact test and we controlled the false discovery rate (FDR) across the tested gene set using the Benjamini–Hochberg (BH). To reduce sparse-count testing, we tested only genes altered in ≥10 HGOS cases total across the full HGOS cohort.

