## Extended Data for "Murine osteosarcoma recapitulates the driver landscape and genomic complexity of osteosarcoma evolution in humans"

### Complex Genomic Rearrangements amplify oncogenes and disrupt tumor suppressor genes in murine model of osteosarcoma

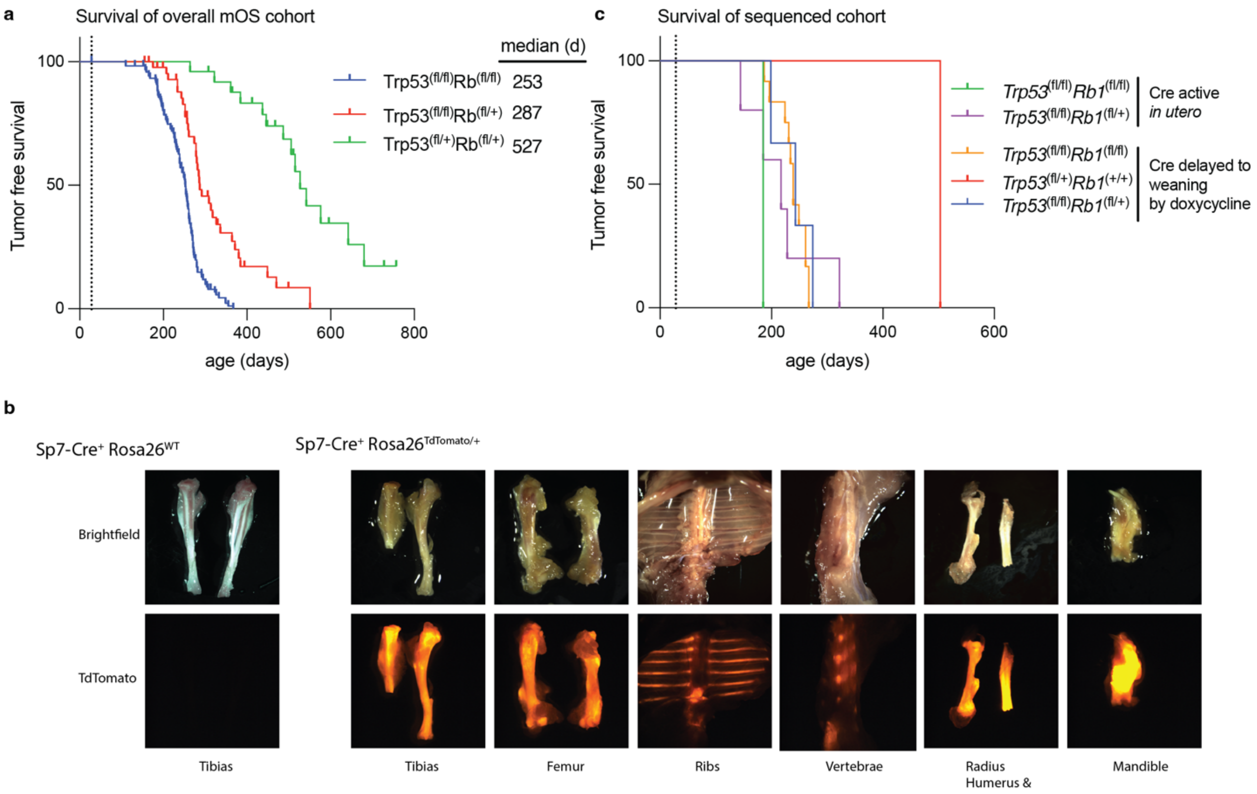


**Extended Data Figure 1 mOS model latency**. a) Tumor free survival of all mOS mice in our cohort (n=234) maintained on doxycycline-food until weaning (d21-28). Mice were monitored weekly for tumor development by exam. Significance assessed by Logrank (Mantel-Cox) test, $p<0.001$. b) Dissection scope images of Sp7-Cre mice bred to Rosa26$^{tdTomato}$ mice. Red fluorescence marks tissues where Cre was active. c) Survival of mice in sequenced cohort (n=24), grouped by genotype and doxycycline treatment.

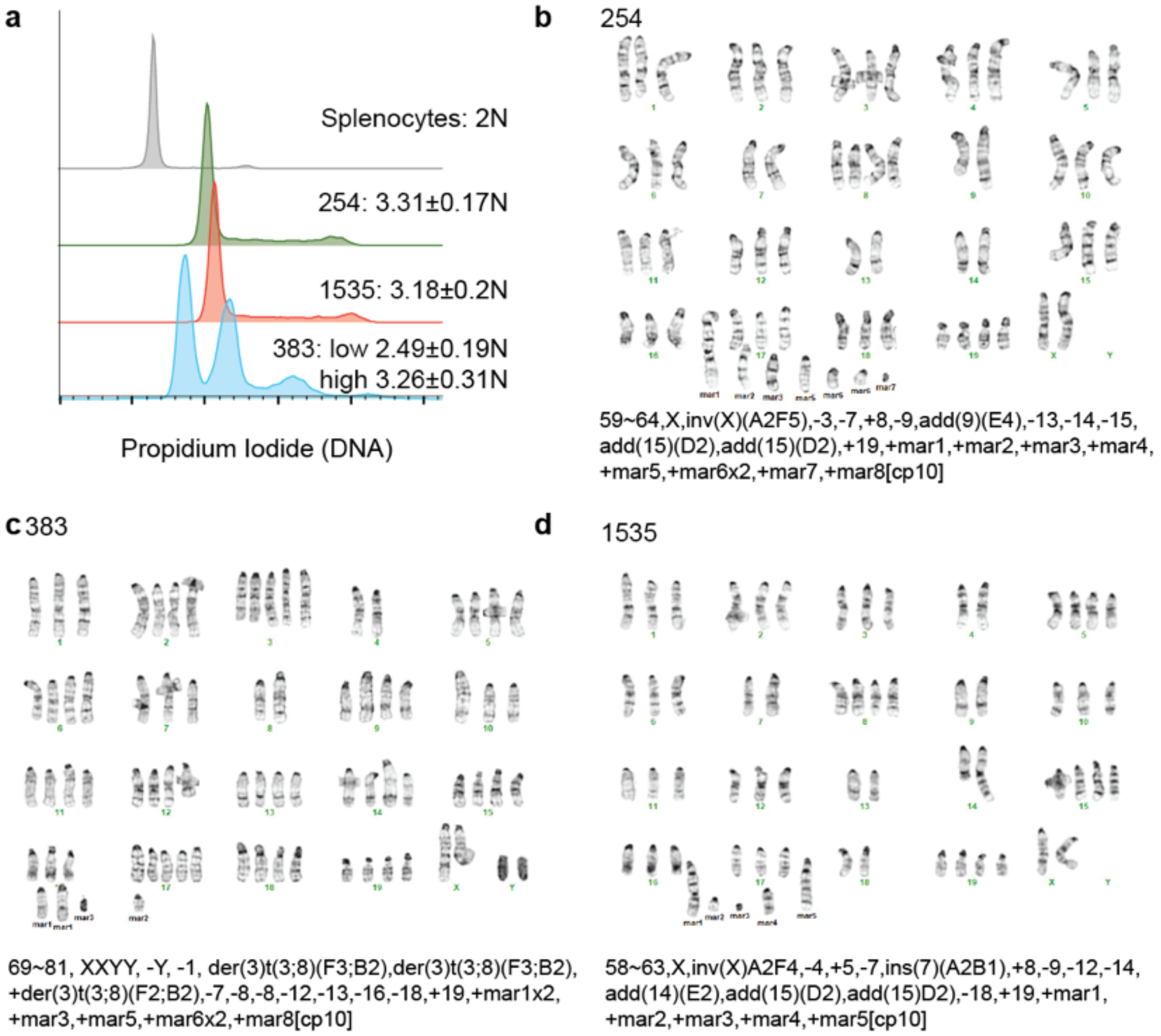


**Extended Data Figure 2 Genome doubling in mOS.** a) DNA content determined by propidium iodine staining relative to healthy murine splenocytes. Calculated copy number determined by relative fluorescence of G1 peak normalized to splenocytes. Reported as mean +/- S.D. from 3 independent experiments. 383 analyzed as mixed population. b-d) Cytogenetic analysis of G-banded metaphase cells from cell lines 254 (b), 383 (c), and 1535 (d). Each image is representative of 10 karyotyped cells.

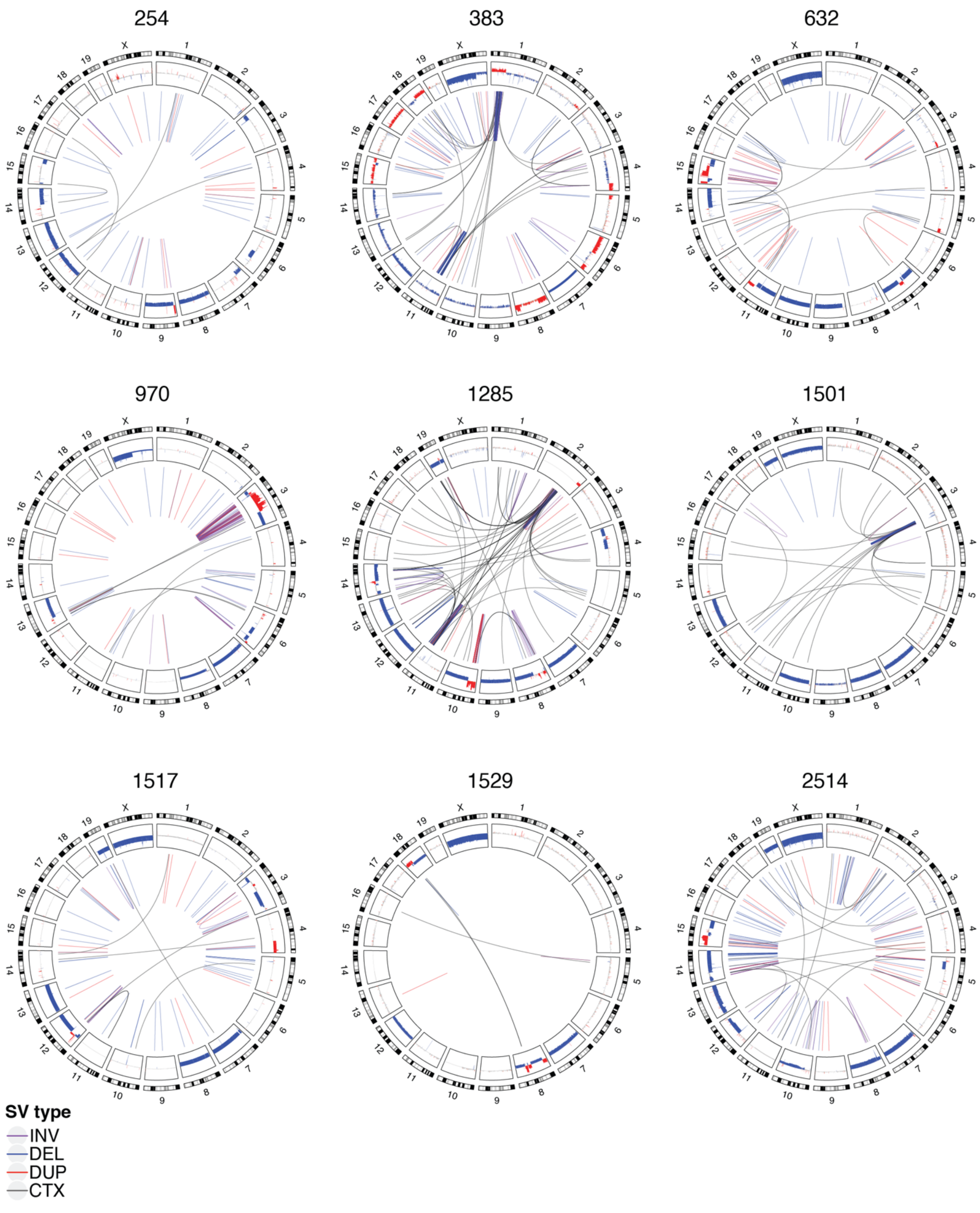


**Extended Data Figure 3 Circos plots of all mOS tumors not in main figures.**

Circos plot tracks depict: (i) chromosome ideograms with Giemsa banding; (ii) SCNAs relative to the tumor baseline ploidy with gains (red) and losses (blue); (iii) SVs depicted by lines and colored by SV type: inversions (purple), deletions (blue), duplications (red) and translocations (black).

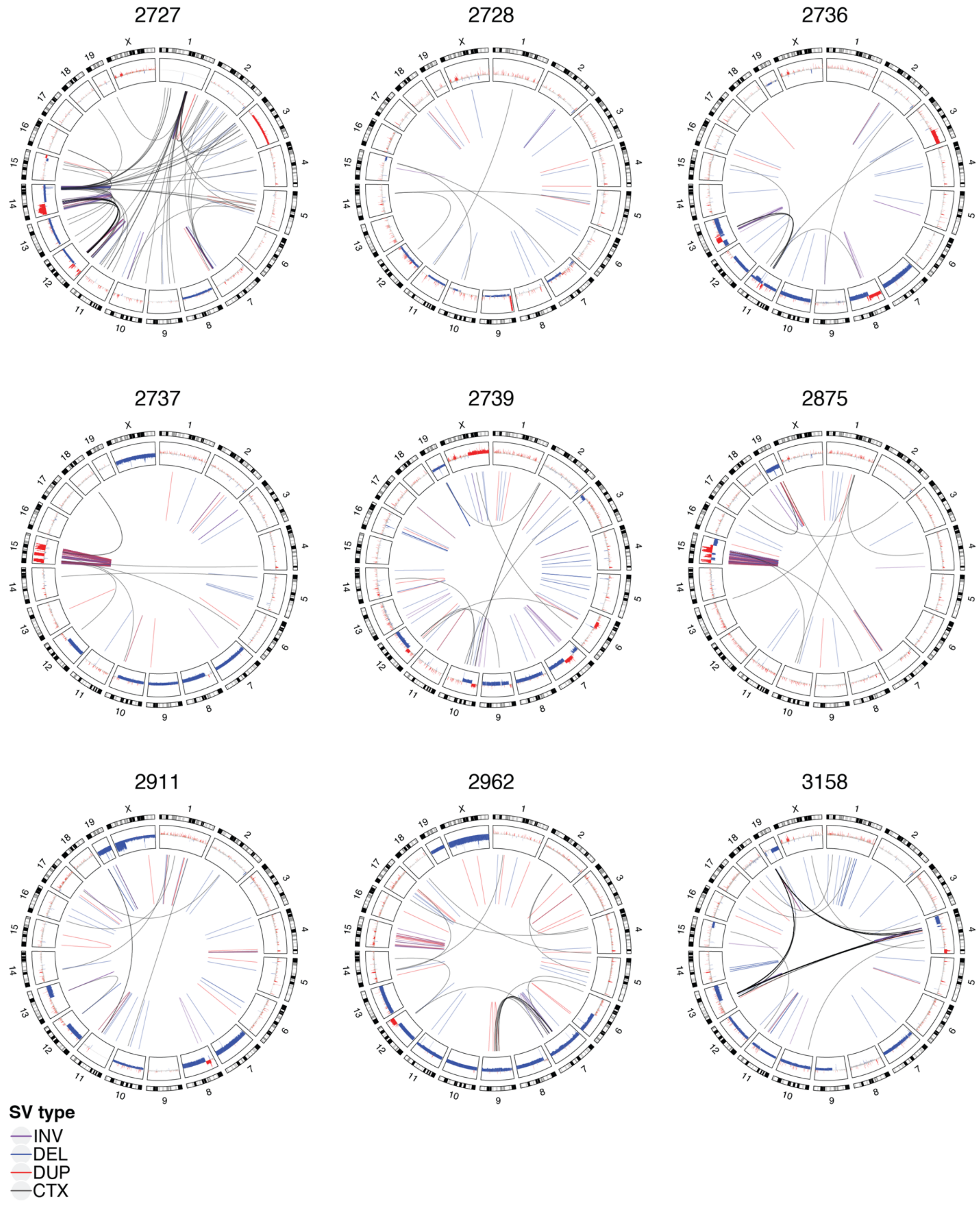


**Extended Data Figure 3 Circos plots of all mOS tumors not in main figures (cont'd).**

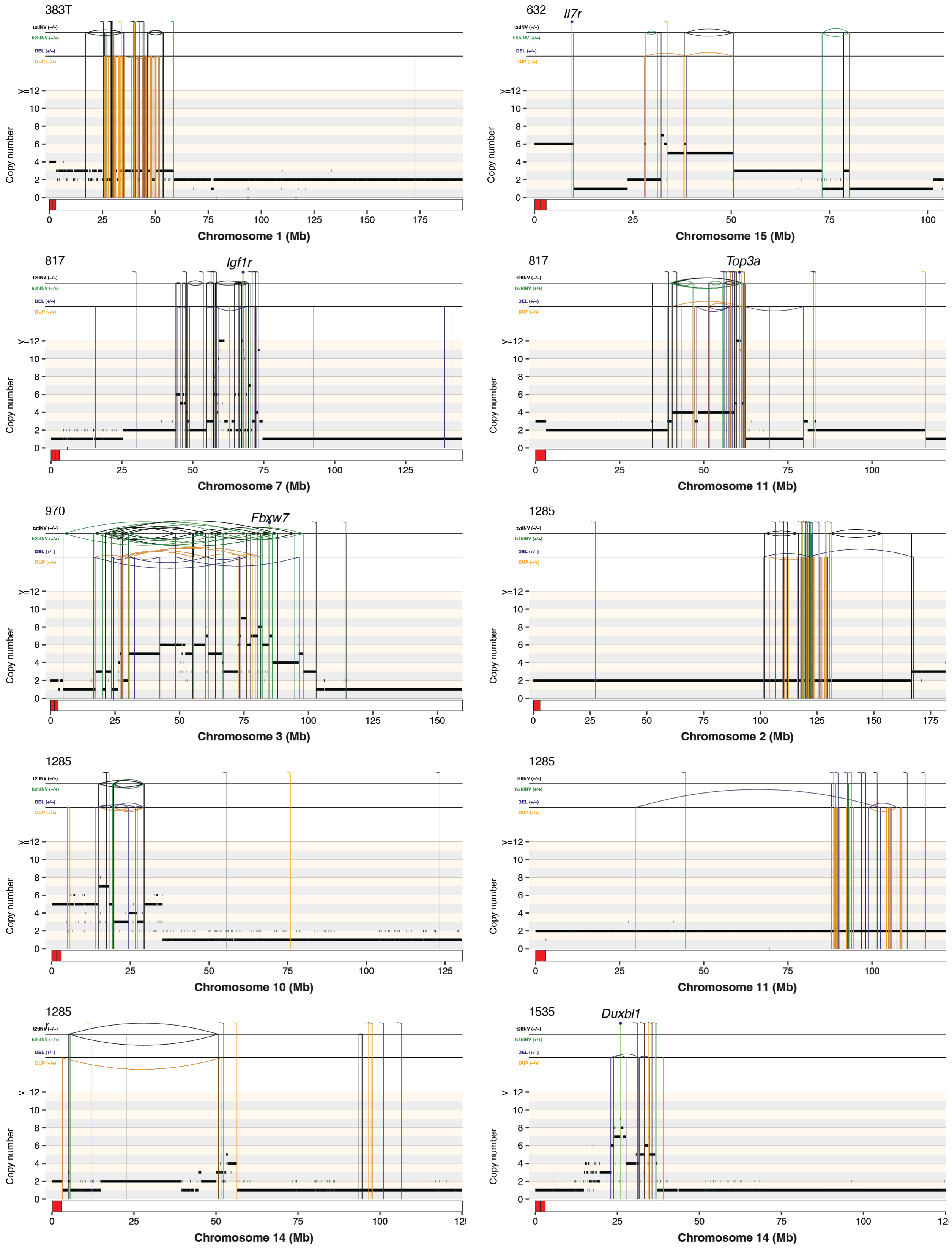


**Extended Data Figure 4 Rearrangement profile plots for CGRs**

Rearrangement profile plots display purity-adjusted rounded copy number values (horizontal lines). SVs are depicted by vertical lines and coloured according to the SV type: DEL (deletion-like) in blue, DUP (duplication-like) in orange, h2hINV (head-to-head inversion) in green, and t2tINV (tail-to-tail inversion) in black.

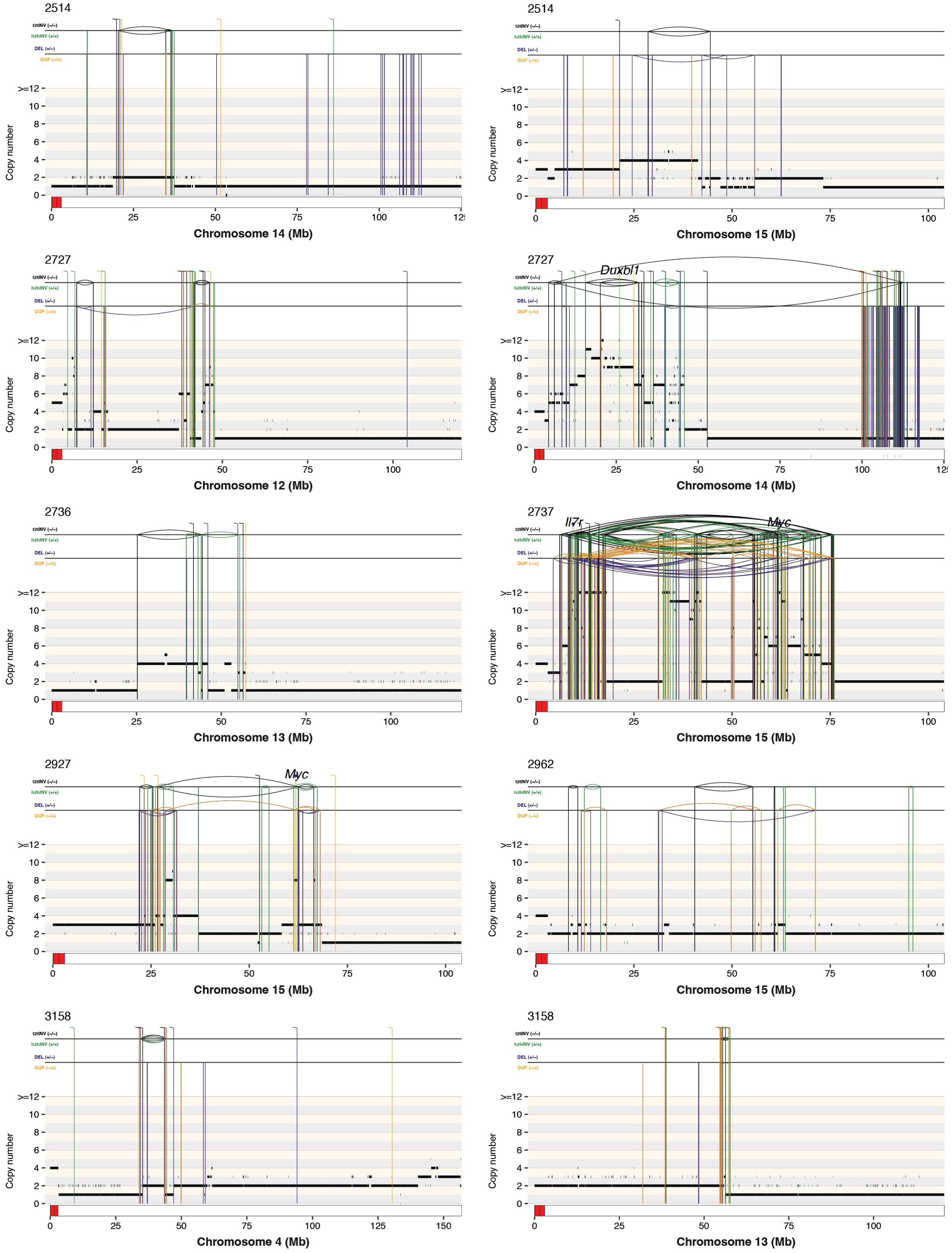


**Extended Data Figure 4 Rearrangement profile plots for CGRs (cont'd)**

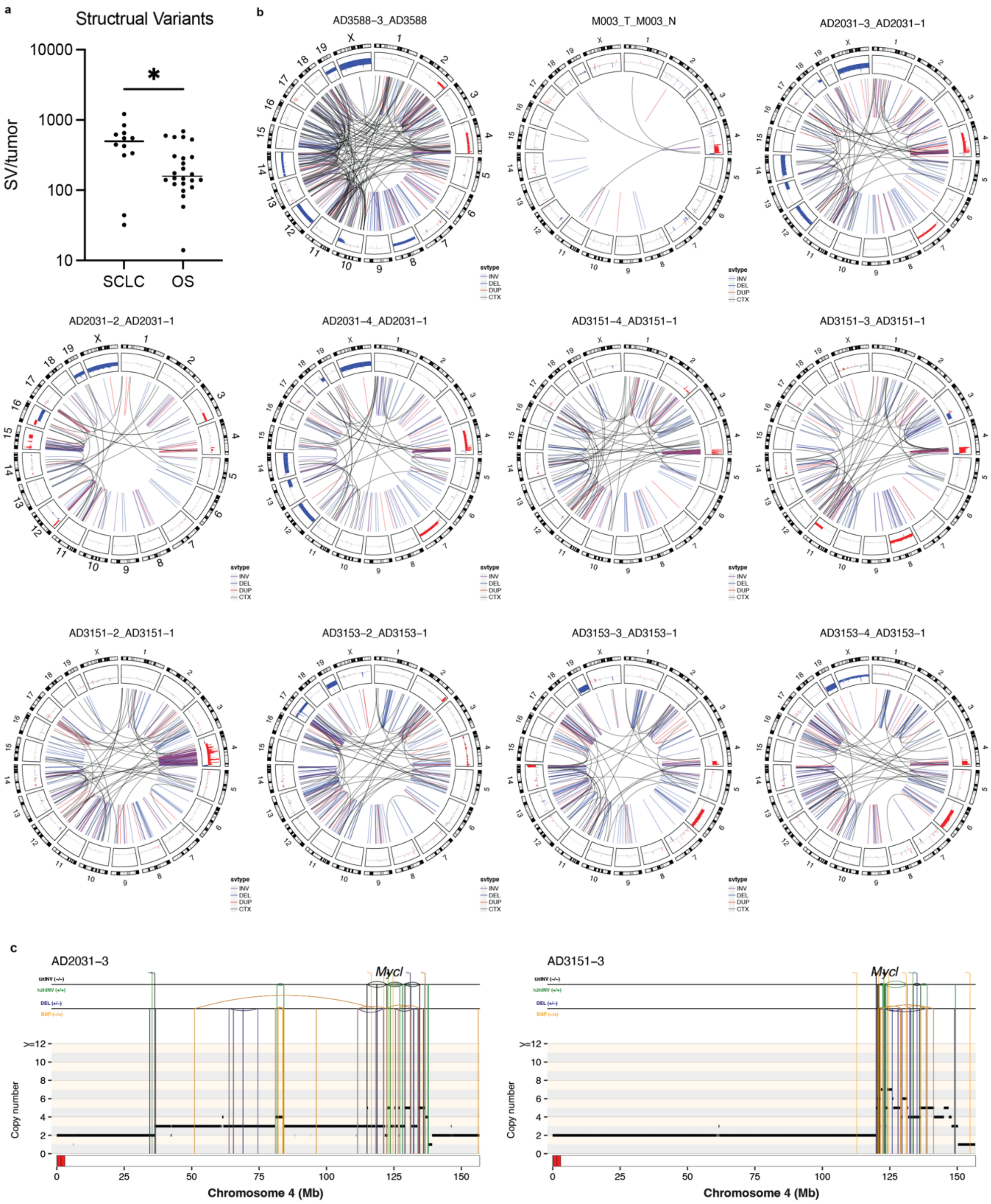


**Extended Data Figure 5 CGRs in murine Small Cell Lung Cancer.** a) Comparison of SV burden in mSCLC to mOS. b) Circos plots of mSCLC tumors. c) Representative rearrangement profile plots of chromosome 4 CGRs overlapping *Mycl*.

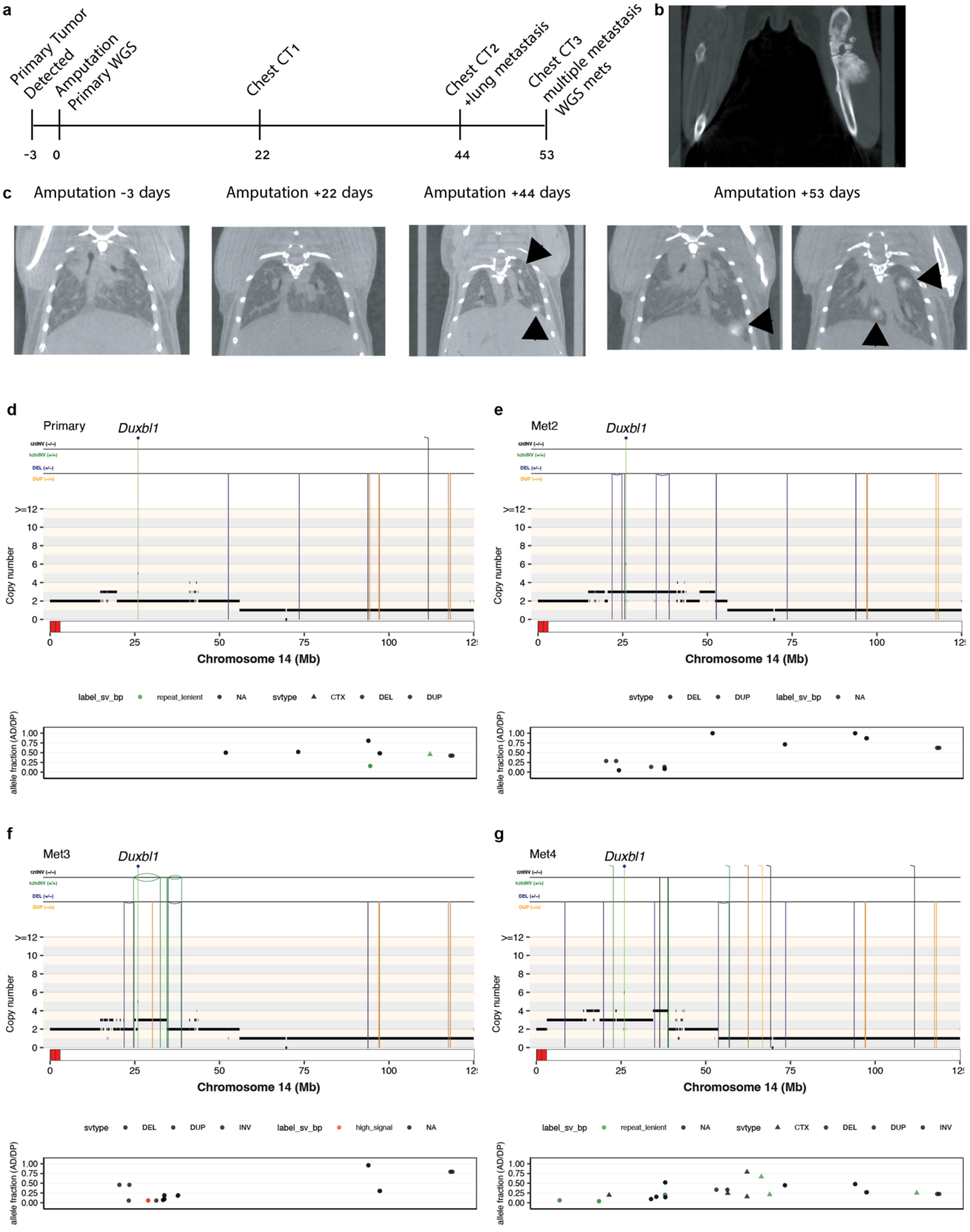


**Extended Data Figure 6. Mouse 727 clinical course and complex rearrangements**. a) Timeline of mouse 727 sample collection and imaging. b) Baseline mCT imaging of left tibia mass prior to amputation. c) Serial chest mCT imaging before and after amputation. Arrowhead marks metastatic lesions. Day +53 display with two coronal slices. d-g) Rearrangement profile plots of CGRs on chromosome 14 with SV variant allele fractions indicated below.

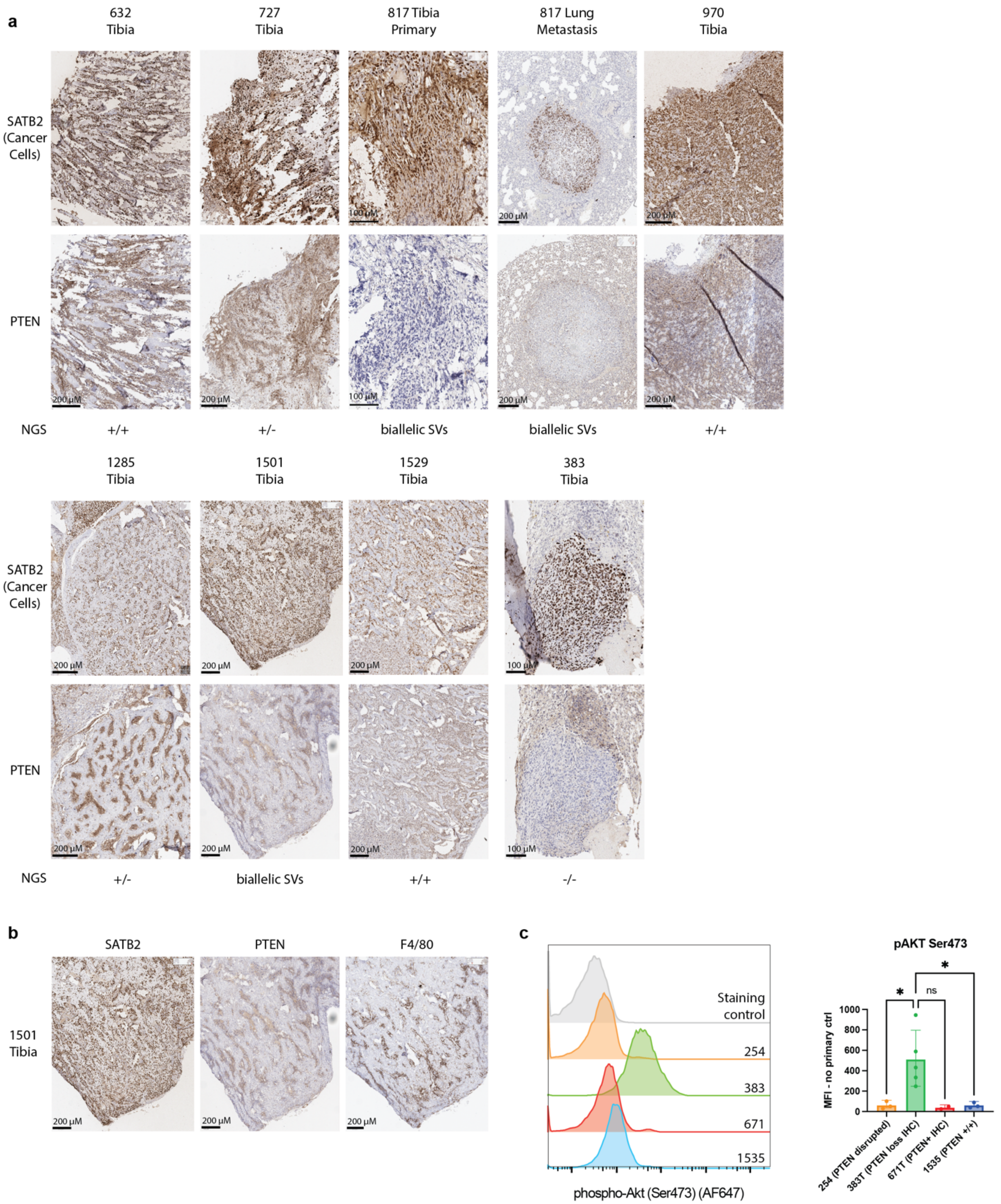


**Extended Data Figure 7 PTEN IHC and Signaling.** a) Immunohistochemistry staining of indicated tumors for SATB2 (expressed in cancer cells) and PTEN. Alterations detected by NGS indicated below. b) Additional staining of sequential sections of tumor 1501 showing PTEN sustaining likely originates from F4/80-positive macrophages and not SATB2-positive cancer cells. c) Histograms of AKT Ser473 phosphorylation as detected by phosphoflow for indicated mOS cell lines, quantified at right over 3 independent experiments. One-way ANOVA with multiple comparisons.

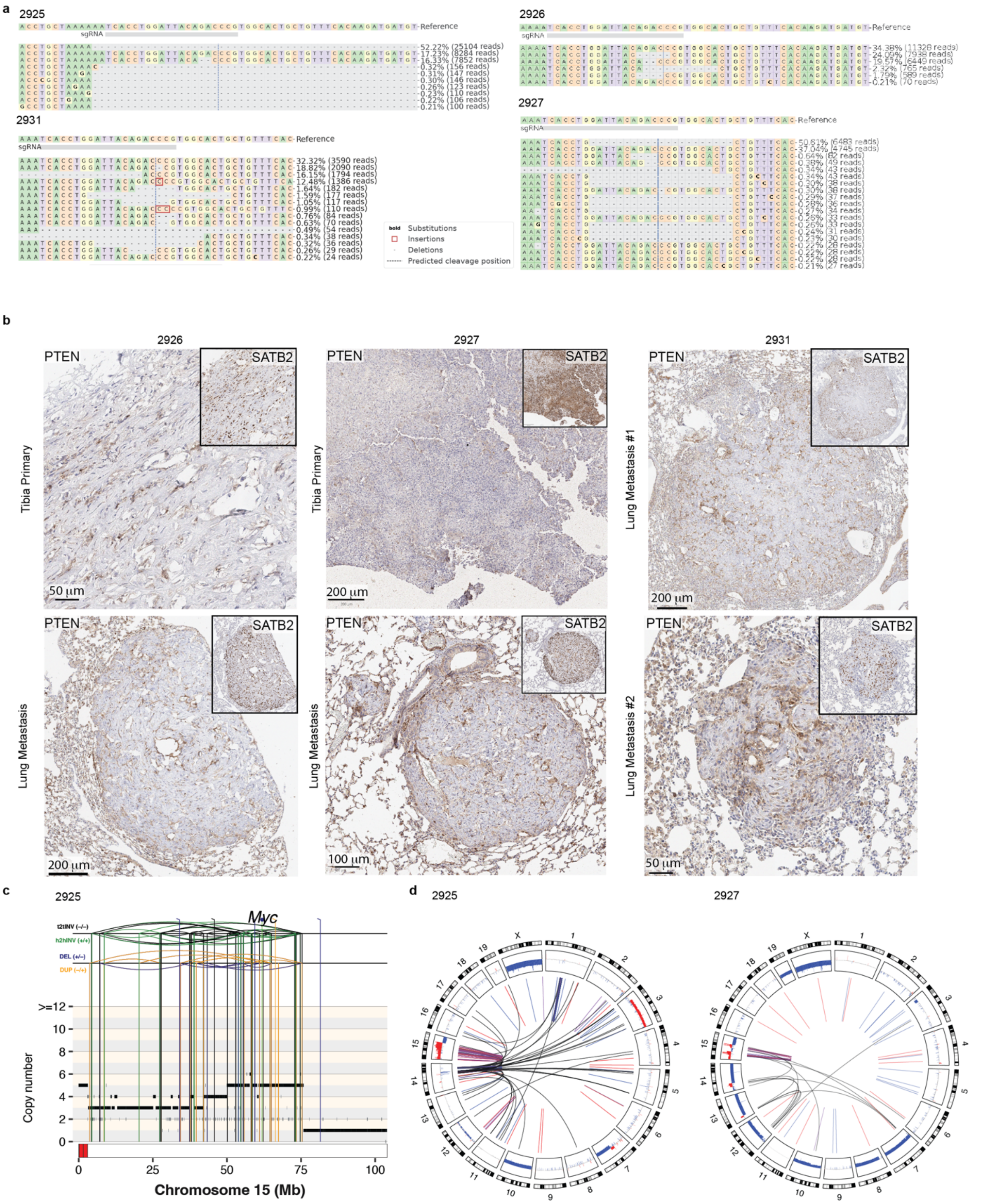

**Extended Data Figure 8 PTEN Editing *In vivo*.** a) Base-level changes following *in vivo* Cas9 editing assessed by Amplicon Sequencing and analyzed with Crispresso2. b) PTEN and SATB2 IHC on edited tumors. c) Rearrangement profile plot of chromosome 15 of PTEN-edited tumor 2925. d) Circos plots from PTEN-edited tumors 2927 and 2925.

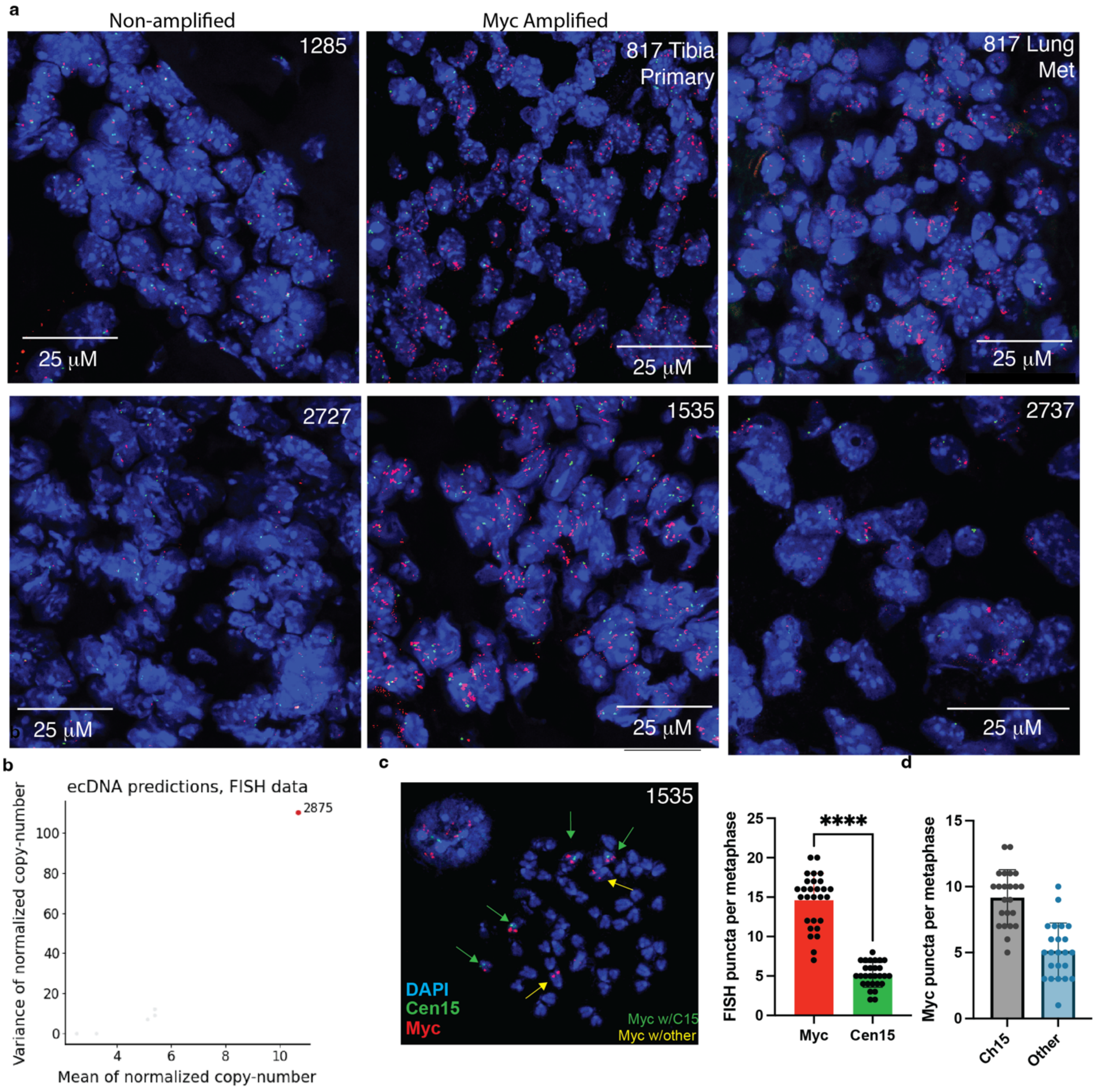


**Extended Data Figure 9. *Myc* FISH FFPE.** A) Representative images of non-amplified and amplified tumor samples, quantified in Fig. 7f. *Myc* probe in red, Centromere 15 in green and DAPI stain blue. B) scAMP ecDNA predictions based on DNA FISH quantification. C) Representative metaphase spread of 1535 cell line, quantified at right (n=23 metaphases, paired t test $p<0.0001$). D) Quantification of *Myc* signals localized to chromosome 15 (Cen15+) and to other chromosomes (n=18 metaphase spreads).

**Supplementary Table 1. Tumor sample characterization and mutation summaries.** Supplementary Table 1.xlsx

**Supplementary Table 2. Gene alterations in murine osteosarcoma.** Supplementary Table 2 Gene Alterations.xlsx

**Supplementary Table 3. Additional Mutations in human *TP53 RB1* double mutant cases.**

| Gene | RB1+TP53 Double Hits | Other Cases | Odds Ratio | p-value | FDR (BH) |
|---|---|---|---|---|---|
| CDKN2A | 0/46 (0.0%) | 21/92 (22.8%) | 0 | 9.98E-05 | ***0.001097*** |
| CCND3 | 0/46 (0.0%) | 14/92 (15.2%) | 0 | 0.004904 | ***0.01798*** |
| CDK4 | 0/46 (0.0%) | 14/92 (15.2%) | 0 | 0.004904 | ***0.01798*** |
| PTEN | 11/46 (23.9%) | 7/92 (7.6%) | 3.816 | 0.01369 | ***0.03764*** |
| CCNE1 | 3/46 (6.5%) | 20/92 (21.7%) | 0.251 | 0.02852 | 0.06274 |
| PIM1 | 1/46 (2.2%) | 12/92 (13.0%) | 0.148 | 0.06 | 0.11 |
| ATRX | 10/46 (21.7%) | 11/92 (12.0%) | 2.045 | 0.1402 | 0.2203 |
| MYC | 3/46 (6.5%) | 10/92 (10.9%) | 0.572 | 0.5435 | 0.7473 |
| RAD21 | 3/46 (6.5%) | 9/92 (9.8%) | 0.643 | 0.7505 | 0.774 |
| NF1 | 5/46 (10.9%) | 8/92 (8.7%) | 1.28 | 0.7597 | 0.774 |
| ERBB4 | 4/46 (8.7%) | 10/92 (10.9%) | 0.781 | 0.774 | 0.774 |

All High Grade OS cases from Valle-Inclan et al.[2] considering genes with 10 or more alteration in the dataset. Analyzed by Fisher’s exact test and false discovery rate controlled by Benjamini-Hochberg procedure.

**Supplementary Table 4: PTEN Editing in mOS Cas9 Mice**

| | ***Background Genotype*** | | | | | | | |
|---|---|---|---|---|---|---|---|---|
| **Case** | ***Trp53*** | ***Rb1*** | ***Rosa26*** | **Outcome** | **Age at tumor (d)** | **Editing Efficiency (WGS purity corrected)** | **Edits** | **PTEN IHC** |
| **2607** | fl/fl | fl/+ | Cas9/+ | Euthanized for poor body condition 81 days w/o tumors | n/a | n/a | n/a | n/a |
| **2612** | fl/fl | fl/+ | Cas9/+ | Mandibular tumor, mScarlet negative | 280 | 0.4% | 4 SNVs, each <0.1% | Pos |
| **2925** | fl/fl | +/+ | Cas9/+ | Left tibia tumor, lung metastasis | 330 | 52.2% (77%) | 58 BP Del | Null |
| **2926** | fl/fl | fl/+ | Cas9/+ | Left tibia tumor, lung metastasis | 376 | 58.56% | 5 BP del (25.6%), 2 BP del (20.7%) | Null |
| **2927** | fl/fl | fl/+ | Cas9/+ | Left Tibia tumor, lung metastasis | 230 | 56.9% (78%) | 21 BP del | Null |
| **2931** | fl/fl | fl/fl | Cas9/+ | Left tibia tumor, lung metastasis | 242 | 63.5% | 1 BP del (18.8%), 22 BP del (16.2%) 1 BP ins (12.48%) | Null |

**Supplementary Table 3**

| Application | Antibody | Manufacturer & Catalog # | Dilution |
|---|---|---|---|
| **IHC** | SATB2 | Abcam AB92446 | 1:250 |
| | PTEN | Cell Signaling Technologies D4.3 #9188 | 1:125 |
| | αRabbit-HRP | Vector Laboratories ImmPRESS HRP Goat Anti-Rabbit IgG MP-7451 | 1 drop |
| **FACS** | phospho-S6 (Ser240/244) | Cell Signaling Technologies D68F8 #5364 | 1:200 |
| | phospho-AKT (Thr308) | Cell Signaling Technologies D25E6 #13038 | 1:1600 |
| | αRabbit-AF647 | Life Technologies A-31573 | 1:500 |
